# The pattern of genetic variability in a core collection of 2,021 cowpea accessions

**DOI:** 10.1101/2023.12.21.572659

**Authors:** Christopher J. Fiscus, Ira A. Herniter, Marimagne Tchamba, Rajneesh Paliwal, María Muñoz-Amatriaín, Philip A. Roberts, Michael Abberton, Oluwafemi Alaba, Timothy J. Close, Olaniyi Oyatomi, Daniel Koenig

## Abstract

Cowpea is a highly drought-adapted leguminous crop with great promise for improving agricultural sustainability and food security. Here, we report analyses derived from array-based genotyping of 2,021 accessions constituting a core subset of the world’s largest cowpea collection, held at the International Institute of Tropical Agriculture (IITA) in Ibadan, Nigeria. We used this dataset to examine genetic variation and population structure in worldwide cowpea. We confirm that the primary pattern of population structure is two geographically defined subpopulations origining in West and East Africa, respectively, and that population structure is associated with shifts in phenotypic distribution. Furthermore, we establish the cowpea core collection as a resource for genome-wide association studies by mapping the genetic basis of several phenotypes, with a focus on seed coat pigmentation patterning and color. We anticipate that the genotyped IITA cowpea core collection will serve as a powerful tool for mapping complex traits, facilitating the acceleration of breeding programs to enhance the resilience of this crop in the face of rapid global climate change.

## Introduction

Enhancing the sustainability of agriculture is essential to feeding the world population, especially in the context of climate change. Among crops, legumes are unique in their ability to improve soil fertility through symbiotic association with rhizobia, which reside within root nodules to facilitate nitrogen fixation (Nap and Bisseling 1990). This quality has made legumes important components of crop rotation and intercropping, both of which help to increase productivity (Reckling *et al*. 2016). In addition, grain legumes are notable for the high protein content of their grains (termed “pulses”), and their amino acid compositions complement those of cereals, making them exceptionally nutritious for both human and animal diets. Productivity gains in grain legumes over the past 50 years have been attributed primarily to expanded planting area rather than to advances in crop improvement as seen in other types of crops, such as cereals (Foyer *et al*. 2016). As such, the development of genetic resources linking genotype to phenotype is crucial to improve grain legumes and to advance crop research and breeding.

Cowpea (*Vigna unguiculata* (L.) Walp.) is a leguminous crop of global importance as a source of both food and fodder. The crop is produced around the world with the majority of contemporary production in West Africa (7.6 of 8.9 million tonnes in 2021) (Food and Agriculture Organization of the United Nations 2022), where it is grown mainly for its protein-rich dry grain. However, other parts of the plant are edible including the leaves, stems, and immature pods (Weng *et al*. 2019; Boukar *et al*. 2019), which, in addition to culinary uses, are used in traditional medicine (Abebe and Alemayehu 2022). Cowpea haulms, stems, and leaves also constitute a nutritious fodder for livestock (Tarawali, S., Singh, B., Peters, M. & Blade, S. 1997). In addition to its ability to grow in nutrient-poor soil, cowpea is also remarkable for having exceptional resilience to drought and high temperatures (Hall 2004), rendering it an ideal candidate for bolstering food system sustainability and for mitigating the repercussions of global climate change.

Linguistic and archaeological evidence suggest that cowpea was domesticated at least 5,000 years ago in Sub-Saharan Africa (Herniter *et al*. 2020), presumably from *V. unguiculata* ssp. *dekindtiana* var. *spontanea* (Pasquet and Padulosi 2012). The domestication region, however, remains a subject of debate with proposed domestication centers in West Africa (Vaillancourt and Weeden 1992; Padulosi and Ng 1997; Coulibaly *et al*. 2002; Ba *et al*. 2004), East Africa (Xiong *et al*. 2018), or in both regions (Huynh *et al*. 2013; Xiong *et al*. 2016; Herniter *et al*. 2020). Previous efforts to study genetic variation in contemporary cowpea germplasm collections have indicated a strong pattern of genetic differentiation between germplasm from West and East Africa (Huynh *et al*. 2013; Xiong *et al*. 2016; Muñoz-Amatriaín *et al*. 2017, 2021; Fatokun *et al*. 2018; Herniter *et al*. 2020), with West Africa as the region of maximum diversity of domesticated cowpea (Fatokun *et al*. 2018; Muñoz-Amatriaín *et al*. 2021).

Although genetic resources for cowpea are underdeveloped compared to cereal crops such as maize, rice, and wheat, cowpea benefits from several available sequenced genomes (Lonardi *et al*. 2019; Liang *et al*. 2023) as well as a high quality genotyping array (Muñoz-Amatriaín *et al*. 2017). Several quantitative trait locus (QTL) mapping populations have been developed to link genotype to phenotype in this species and have been useful in identifying QTL for diverse traits such as flowering time (Andargie *et al*. 2013; Huynh *et al*. 2018; Lo *et al*. 2018; Olatoye *et al*. 2019), disease resistance (Huynh *et al*. 2016; Santos *et al*. 2018), seed coat color and patterning (Herniter *et al*. 2018, 2019), perenniality (Lo *et al*. 2020), and several domestication-related traits such as seed shattering and yield (Lo *et al*. 2018). Additionally, several genome-wide association studies (GWAS) have been conducted to discover SNPs associated with flowering time (Paudel *et al*. 2021; Muñoz-Amatriaín *et al*. 2021), disease resistance (Kpoviessi *et al*. 2022; Dong *et al*. 2022), seed size (Lo *et al*. 2019), seed protein content (Chen *et al*. 2023), drought response (Ravelombola *et al*. 2021; Wu *et al*. 2021), pod shattering (Lo *et al*. 2021), and salt tolerance (Ravelombola *et al*. 2022). Nevertheless, the majority of QTL mapping studies in cowpea have been limited in their ability to fine-map associated loci due to constrained sample sizes or low molecular marker densities.

Here, we report the genetic characterization of a core collection of cowpea held at the International Institute of Tropical Agriculture (IITA), a CGIAR research center in Ibadan, Nigeria. The IITA cowpea collection represents the most comprehensive collection of worldwide cowpea germplasm, currently consisting of 16,460 accessions collected from 91 countries (March 2022) (IITA 2023). The bulk of the collection (~12,000 lines) consists of locally adapted landraces (hereafter traditional cultivars). The core subset of the collection was selected to represent the diversity of geographic origin and agrobotanical traits present among accessions in the full collection (Mahalakshmi *et al*. 2007) and has been subsequently refined to include 2,082 accessions. To study genetic variation in the Core Collection, we made use of previously generated SNP data (Close *et al*. 2023) consisting of 2,021 accessions genotyped at nearly 50,000 polymorphic sites using the Illumina Cowpea iSelect Consortium Array (Muñoz-Amatriaín *et al*. 2017). We then leveraged this dataset to characterize the pattern of population structure in global cowpea and demonstrated the utility of this resource for genome-wide association studies by dissecting the genetic basis of variation in seed pigmentation.

## Results

### Genetic diversity of the IITA Cowpea Core Collection

Genomic DNA was extracted from 2,021 accessions from the IITA Cowpea Core Collection and scored at nearly 50,000 polymorphic sites using the Illumina Cowpea iSelect Consortium Array (Close *et al*. 2023). Following removal of sites and samples with missing data and excess heterozygosity, the dataset comprised genotypes from 1,991 accessions collected from 86 countries across 6 continents (Figure 1A). All major cowpea producing regions were represented in the dataset with 1,547 accessions from Africa, 207 from Asia, 110 from North America, 68 from Latin America, 43 from Europe and 14 from Oceania. The accessions mostly consisted of 1,590 traditional cultivars, with an additional 206 breeding lines mostly from Nigeria and the United States of America. Additionally, there were 57 uncultivated accessions, presumed to be wild or weedy.

**Figure 1.**
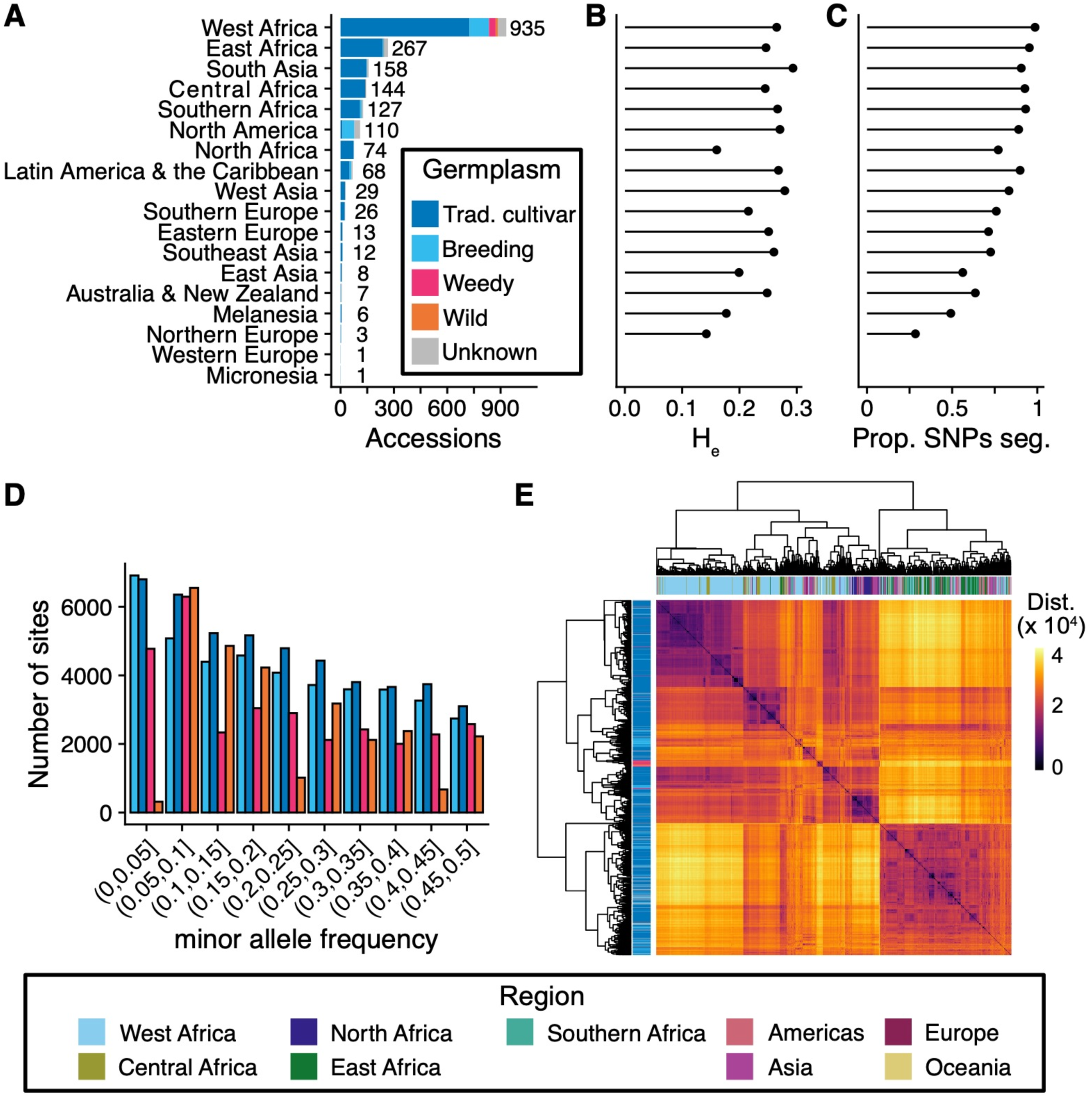
Genetic diversity of the IITA Core Cowpea Collection. A) Region of origin and germplasm status for genotyped accessions. Labels beside each bar indicate the number of accessions per region. B) Average expected heterozygosity per region of origin calculated using all sites. C) Proportion of SNPs segregating per region. D) Folded allele frequency spectrum by germplasm status. E) Heatmap of pairwise genetic distances. Row labels indicate germplasm status and column labels indicate region of origin.

Pairwise genetic distances calculated between accessions across filtered sites revealed many nearly identical accessions in the Core Collection (Figure S1). In total, we identified 130 clusters (comprising 400 accessions) of nearly identical accessions by applying hierarchical clustering to the genetic distance matrix (Table S1, static tree cut threshold of 500 corresponded to differences at ~0.5% or fewer sites). The vast majority of clusters consisted of either two or three accessions (79 and 25 clusters, respectively), while 26 clusters consisted of 4 or more accessions (max cluster size 15). Most clusters (72 / 130) contained accessions collected from different countries with 40 clusters consisting of accessions from countries on different continents, 19 clusters consisting of accessions collected from adjacent countries within Africa, 12 clusters consisting of accessions from non-adjacent countries within Africa, and 1 cluster with accessions from adjacent countries in South America. The geographic distribution of accessions within clusters suggests that accessions were dispersed between continents by human activity and then resampled during the construction of the Core Collection. Of the 130 clusters, 86 consisted of accessions of the same germplasm status (e.g. traditional cultivars only), 20 clusters contained both traditional cultivars and breeding lines, 12 clusters had traditional cultivars and accessions of unknown status, 3 clusters contained breeding material and accessions of unknown status, and 2 clusters contained wild or weedy accessions and traditional cultivars. We randomly selected one accession from each of the 130 clusters for subsequent analysis.

The final dataset consisted of 1,722 accessions genotyped for 48,004 biallelic SNP markers. The included accessions reflect the geographic diversity of the IITA Cowpea Collection, with lines originating from 85 countries and include ~83% (1,722 / 2,081) of accessions from the IITA Cowpea Core Collection (Mahalakshmi *et al*. 2007) and 86% (324 / 376) of accessions included in the IITA Cowpea Mini-Core Collection (Fatokun *et al*. 2018). The analyzed samples encompassed 1,394 traditional cultivars, 171 breeding and research lines, 32 weedy accessions, 17 wild accessions, and 110 lines with unknown germplasm status. While the traditional cultivars constitute a global collection, the breeding and research material was predominately collected from Africa and North America (98 and 51 accessions, respectively). The presumed wild accessions were exclusively collected from Africa, while the presumed weedy accessions originated solely from West Africa.

Crop dispersal and improvement can lead to genome-wide depletion of genetic diversity. We sought to determine how these processes impacted cowpea by estimating average expected heterozygosity (H_e_) within the major geographic regions represented in our dataset. H_e_ ranged from 0.14 in Northern Europe to 0.29 in South Asia among regions that contributed 3 or more accessions (Figure 1B, Table S2). Within Africa, H_e_ was similar for all regions (H_e_ = 0.25-0.27) except for North Africa, which had a substantial reduction in diversity (H_e_ = 0.15). Outside of Africa, H_e_ varied considerably, with higher values exceeding those observed in Africa in South Asia, West Asia, North America, and Latin America. In contrast, all regions of Europe and Oceania had reduced H_e_ compared to Sub-Saharan Africa.

Although H_e_ provides a summary of genetic variation within a population, the measure is influenced by the number of segregating alleles and their frequencies. The number of segregating sites per geographic region ranged from 13,660 in Northern Europe to 47,361 in West Africa (Figure 1C, Table S2). The number of segregating sites per region was greatest in Sub-Saharan Africa, with a loss of over 10% of segregating sites observed in all other regions except for South Asia. The proportion of segregating sites was especially low in Oceania and East Asia, though these regions were not well sampled in the IITA core.

Taken together, our analyses are consistent with the hypothesis that West Africa is the center of domesticated cowpea genetic diversity with the largest number of segregating sites observed in this region. We observed a reduction of genetic diversity in all regions out of Africa except South Asia, which is consistent with founder effects upon dispersal. The elevation of H_e_ in South Asia is largely due to the majority of alleles segregating at intermediate frequencies (median minor allele frequency among segregating alleles = 0.25). The reduction of both H_e_ and the proportion of segregating sites in North Africa suggests strong founder effect with low genetic diversity among the founding lines. We also found evidence that cowpea improvement has reduced genetic diversity, with breeding lines having an enrichment for rare alleles when compared to traditional cultivars (Figure 1D). Of the 47,660 SNPs segregating amongst traditional cultivars, only 46,298 segregated in the breeding material.

### Patterns of population structure

After removal of nearly identical samples, we observed a multimodal distribution of pairwise genetic distances between accessions suggesting that strong population structure might exist in the core collection. Hierarchical clustering of accessions by genetic distance linked population structure to geographic origin and germplasm status (Figure 1E). The primary genetic pattern in the dataset consisted of two large clusters: one cluster composed of accessions from Eastern and Southern Africa and a second cluster encompassing accessions from Northern, Central, and Western Africa. Germplasm collected from outside Africa is interspersed within the two main clusters, consistent with the aforementioned pattern of migration out of Africa likely facilitated by human activity. We also observed that many wild and weedy accessions clustered together within the larger cluster. It remains uncertain whether this observation is a result of ascertainment bias of our genotyping approach or if accessions designated as wild or weedy are actually feral accessions that have escaped cultivation.

To examine population structure in more detail, we next estimated the ancestries of each accession using ADMIXTURE (Alexander *et al*. 2009; Alexander and Lange 2011) (Figure 2A). Evanno’s ΔK method (Evanno *et al*. 2005) suggested an optimum of two subpopulation clusters (Figure S2). We assigned accessions to the respective cluster that contributed 70% or more ancestry, resulting in 614 accessions in cluster 1, 680 accessions in cluster 2, and 428 accessions unassigned to a cluster (Table S1). Accessions were non-randomly distributed between clusters for 13 of the 17 geographic regions (Chi-squared tests, p < 0.05, Table S3). Cluster 1 contains the majority of accessions from East Africa (90%), Latin America and the Caribbean (75%), North America (71%), Southern Africa (72%), and Australia and New Zealand (67%). In contrast, cluster 2 consists of the majority of accessions from North Africa (92%), West Africa (59%), and Central Africa (52%). The majority of lines from East Asia (86%), Melanesia (83%), and Southeast Asia (82%) were unassigned to a cluster. Germplasm from Eastern Europe, Micronesia, Southern Europe, and West Asia did not have significant enrichment for belonging to a particular cluster. Given the strong partitioning of accessions by geographic subregion within Africa, we hereafter refer to cluster 1 as the ESA (Eastern and Southern Africa) cluster and cluster 2 as the WNCA (West, North, and Central Africa) cluster. The two clusters have significant genetic differentiation (F_st_ = 0.292) and the ESA cluster has a slightly higher level of genetic diversity compared to the WNCA cluster (H_e_ = 0.25 for ESA, 0.22 for WNCA, 0.28 for unassigned).

**Figure 2.**
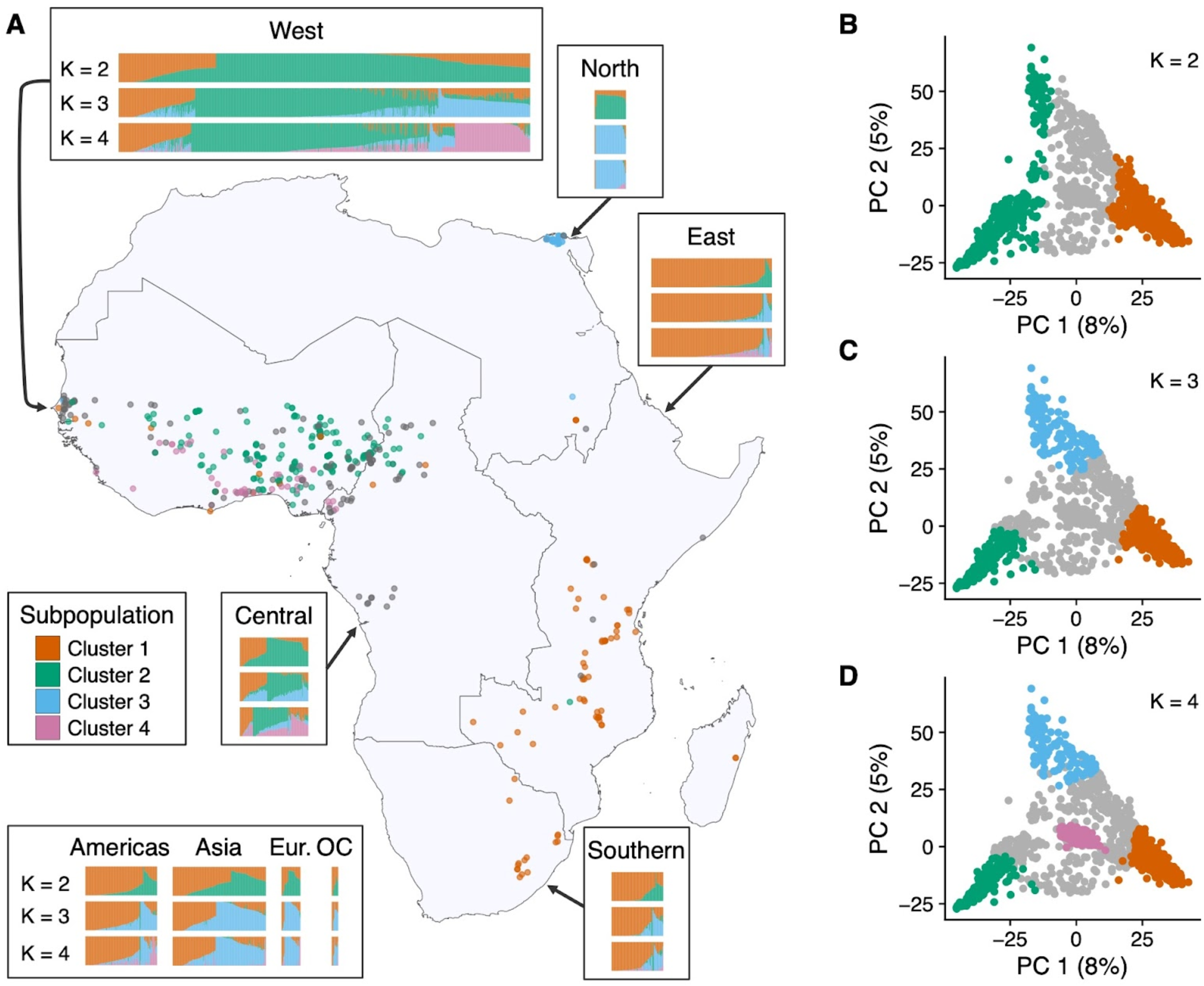
Population structure in global cowpea germplasm. A) Inferred ancestry for all genotyped individuals from the core collection and map of collection sites for 545 traditional cultivars with known collection sites in Africa. Accessions are grouped by subregion according to the United Nations geoscheme and each accession with a known collection site is colored according to subpopulation assignment when modeling K = 4 subpopulations. Accompanying barplots specify the proportions of ancestry attributed to each subpopulation per individual when modeling K = 2, K = 3, and K = 4 subpopulations, respectively. Eur. = Europe, OC = Oceania. B - D) Principal components analysis of allele frequencies with accessions colored by subpopulation for K = 2 - 4.

To explore the possibility of population substructure in West Africa, we considered additional subpopulations. Considering K = 3 subpopulations reassigned 124 accessions from the WNCA cluster and 82 previously unassigned accessions to the new cluster 3. Additionally, 78 and 146 accessions from the ESA and WNCA clusters, respectively, were considered to be unassigned at K = 3. Cluster 3 consisted of nearly all accessions from North Africa in proximity to the Nile River Delta (57 / 62) and the majority from Southern Europe (20 / 25), East Asia (5 / 7), and Eastern Europe (6 / 10) (Figure 2A). We also observed that the ancestry previously attributed to the WNCA cluster in accessions found outside of Africa was reassigned to cluster 3. Cluster 3 accessions were nearly exclusively collected in arid climates (Köppen climate classification: BWh/BWk), while other clusters are found to be mostly confined to wet tropical climates (Figure S3), hinting at the possibility that cluster 3 ancestry includes alleles locally adapted to warmer, drier environments.

**Figure 3.**
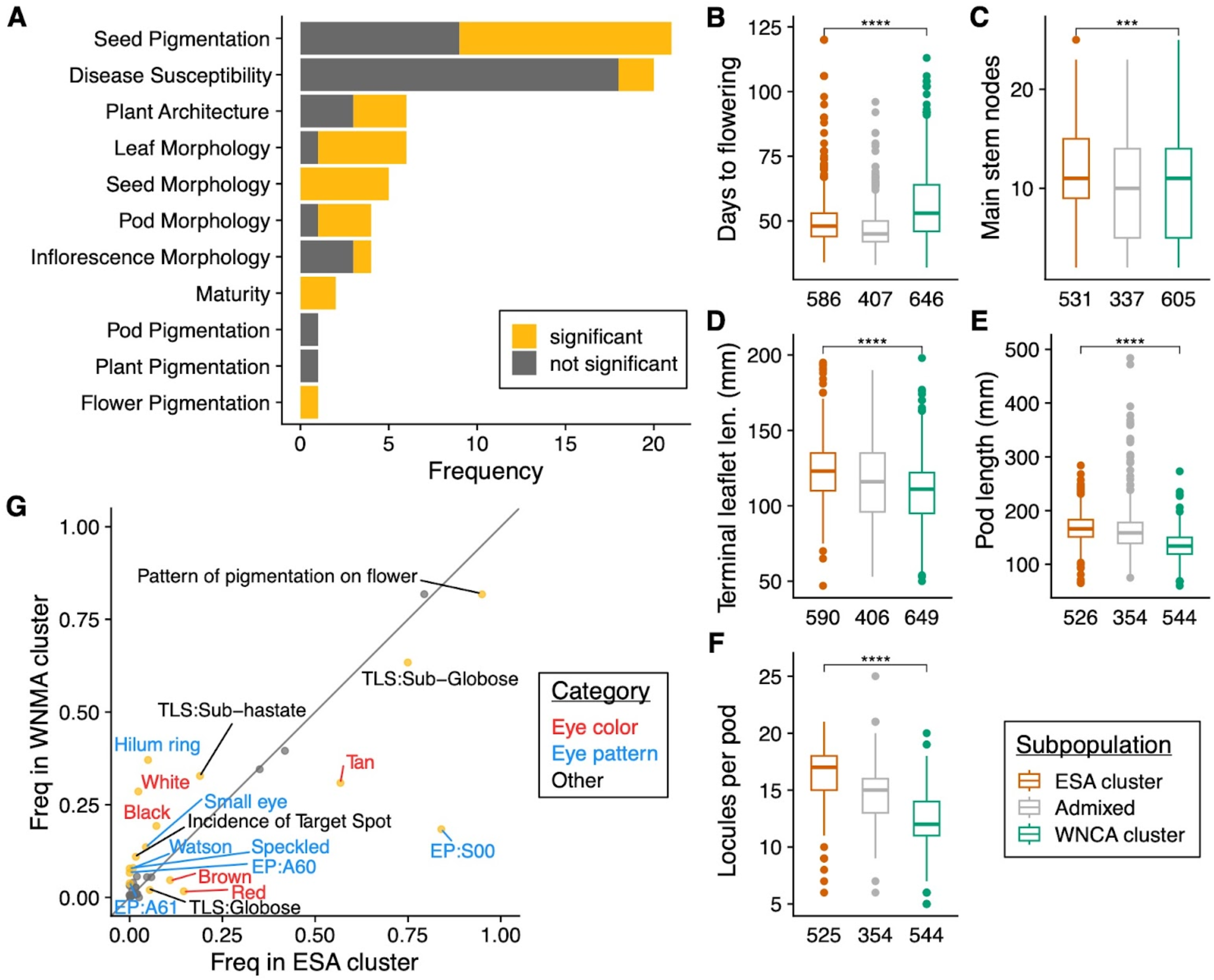
Phenotypic distributions in subpopulation clusters. A) Number of phenotypes by category, colored by the number of phenotypes with statistically significant difference between the ESA and WNCA clusters. B-F) Tukey’s boxplots of representative phenotype distributions by subpopulation cluster. Numbers below each boxplot denote N. Additional phenotypes are presented in Figure S6. G) Qualitative trait frequencies in the two subpopulation clusters. Grey reference line indicates equal frequency in both clusters. Points are colored according to whether there is a significant difference in trait frequency (goldenrod) between the two subpopulation clusters. Labels correspond to phenotype names and are colored by category. Abbreviations: EP-eye pattern, TLS-terminal leaf shape.

Modeling population structure with an additional subpopulation (K = 4) grouped 148 accessions previously unassigned to a cluster when considering fewer subpopulations to cluster 4 (Figure 2A,D). Cluster 4 consisted of 133 accessions from West Africa, 7 accessions from either Central or East Africa, and 8 accessions collected in other world regions. The West African cluster 4 accessions were not associated with a particular landform such as a river or mountain range and, like the majority of accessions in the WNCA cluster, originated from tropical savanna climates (Figure S3). This group of accessions likely represents early-flowering traditional varieties that were cultivated alongside later maturing lines as a precaution against drought (Mortimore *et al*. 1997).

A principal component analysis (PCA) of allele frequencies was largely consistent with the population structure modeling. The first 10 PCs explain 27% of the variance in the dataset, with nearly half of the variance (12.8%) explained by the first 2 PCs alone (Figure S4). Overall, the distribution of individuals along the first two PCs was continuous, suggesting frequent admixture between population groups (Figure 2B-D). PC 1 separates the ESA cluster from the WNCA cluster (Figure 2B-D), while PC 2 separates cluster 3 from the other clusters (Figure 2C). Cluster 4 is not resolved along the first two PCs and is instead resolved on PC 3 (Figure 2D, Figure S5).

**Figure 4.**
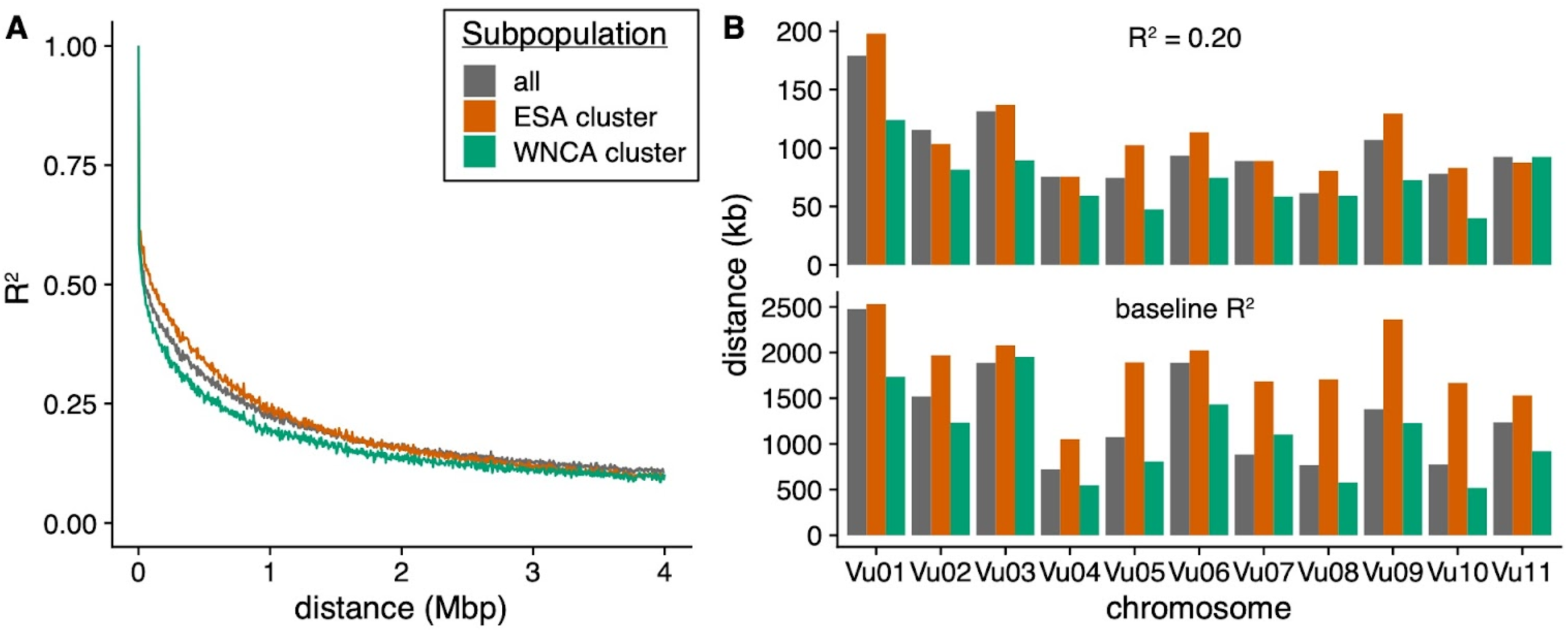
Linkage disequilibrium in the cowpea genome. A) Mean pairwise linkage disequilibrium (LD) between SNPs in 500 bp windows. B) LD per chromosome. Minimum distance to R^2^ of 0.20 (top) and to chromosome-specific background level of LD (bottom).

**Figure 5.**
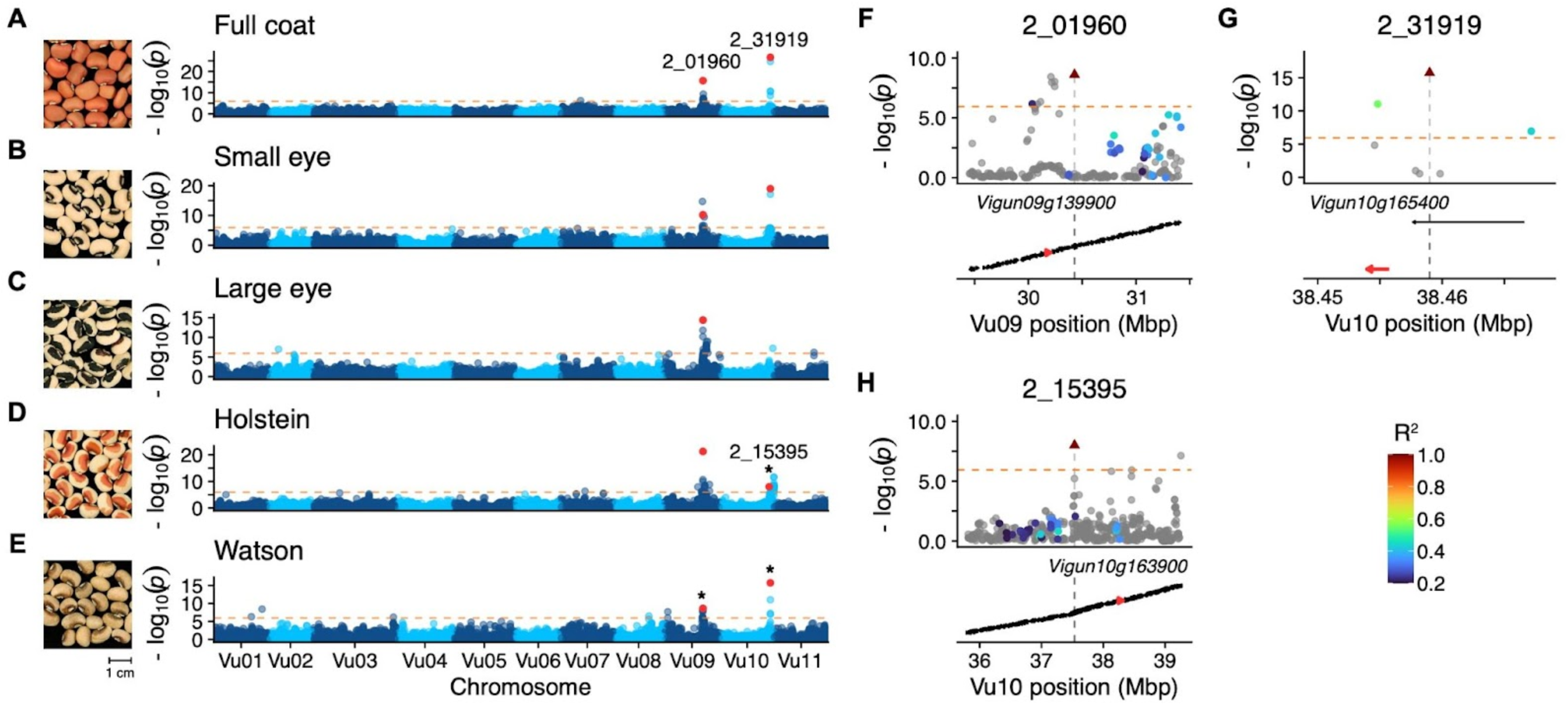
The genetic basis of seed coat pigmentation pattern. A-E) Manhattan plots from genome-wide association mapping of seed coat pigmentation pattern phenotypes. Orange line indicates Bonferroni corrected significance threshold. Tag SNPs are highlighted in red and labeled. Tag SNP 2_15395 is unique to Holstein, highlighted SNP on Vu10 in Watson and Full coat correspond to 2_31919. Photographs from UCR Cowpea Minicore as follows: Self colored (TVu-3842/MinicoreUCR_247), Small eye/Eye 1 (TVu-16278/MinicoreUCR_197), Large eye/Eye 2 (TVu-8775/MinicoreUCR_318), Holstein (TVu-7625/MinicoreUCR_290), Watson(TVu-2933/MinicoreUCR_232). F-H) Local linkage equilibrium for tag SNPs (triangle) for peaks indicated by * in A-E. Candidate genes are indicated in red and labeled.

ESA ancestry was also overrepresented in the modern breeding material present in the IITA core though a large proportion of accessions remained unassigned (Table S4), perhaps the result of controlled crosses between accessions from the two clusters. Consistently, increasing K did not reduce the number of unassigned accessions, pointing to true admixture rather than cryptic population structure in the sample. In addition, we found limited evidence for contribution from clusters 3 and 4 in the breeding germplasm.

Overall, our population structure analyses confirm that global cowpea is characterized by genetic differentiation between germplasm from Western versus Eastern and Southern Africa (Huynh *et al*. 2013; Xiong *et al*. 2016; Muñoz-Amatriaín *et al*. 2017, 2021; Fatokun *et al*. 2018; Herniter *et al*. 2020). Further, our results suggest that germplasm derived from Northern, Eastern, and Southern but not Western Africa represent the majority of ancestry in lines distributed to other parts of the world. Breeding material appears to be enriched for ancestry from the ESA cluster, potentially neglecting climate-adapted alleles from other clusters.

### Phenotypic distributions of subpopulations

Our analyses of population structure in global cowpea indicate that cowpea germplasm is broadly genetically differentiated into two germplasm groups defined by geography (i.e. the ESA and WNCA clusters). To understand the degree to which population structure is linked to phenotype, we leveraged the IITA Cowpea Characterization and Cowpea Evaluation datasets (IITA 2023) to compare the distributions of 34 quantitative phenotypes and the frequencies of 37 qualitative phenotypes between the two subpopulation clusters (Materials and Methods, Table S5, Table S6). As a caveat, the phenotyping data we analyze here was collected through IITA’s cowpea breeding program through numerous field trials conducted across both time (1967-present) and location. Therefore, the data represent reference phenotype values used for breeding efforts and do not correspond to values measured as part of a single experiment.

We identified 17 quantitative phenotypes with significantly different distributions between the WNCA and ESA subpopulation clusters (Wilcoxon test Bonferroni-corrected p < 0.05, Figure 3A, Table S7). Overall, the majority of differences were found in life history and morphological traits and not in disease susceptibility traits. Accessions in the ESA cluster generally matured earlier (Figure 3B), were more branched (Figure 3C), had larger lateral organs (Figure 3D-E), and produced more seeds that were generally smaller in size compared to the WNCA cluster (Figure 3F). Additionally, accessions in the ESA cluster were more likely to have smooth compared to rough testa texture, raceme in all layers of the canopy rather than in the upper leaf layers, intermediate plant growth habit with lower branches touching the ground, more twining, and higher susceptibility to anthracnose compared to the WNCA cluster (Figure S6).

**Figure 6.**
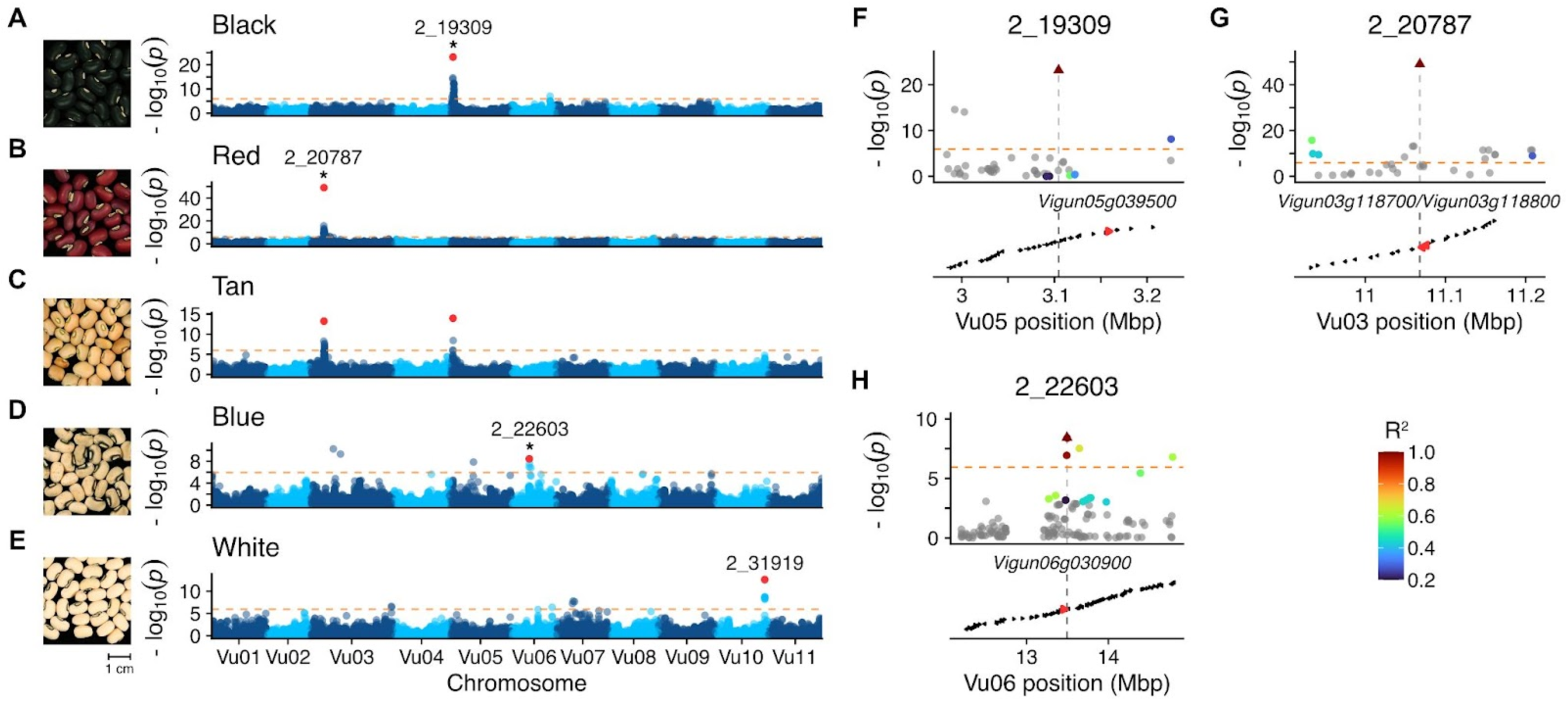
The genetic basis of seed coat color. A-E) Manhattan plots from genome-wide association mapping of seed color phenotypes. Orange line indicates Bonferroni corrected significance threshold. Tag SNPs are highlighted in red and labeled. Photographs from UCR Cowpea Minicore as follows: Black (TVu-13968/MinicoreUCR_139), Red (TVu-15426/MinicoreUCR_173), Tan (TVu-1036/MinicoreUCR_108), Blue (TVu-4669/MinicoreUCR_262),White (TVu-14401/MinicoreUCR_153). F-H) Local linkage equilibrium for tag SNPs (triangle) for peaks indicated by * in A-E. Candidate genes are indicated in red and labeled.

We also observed differences in the incidence of qualitative traits between the ESA and WNCA subpopulation clusters. In total, there were 17 traits with differential frequency between the two clusters (Fisher’s Exact Test Bonferroni-corrected p < 0.05, Figure 2G, Table S8). Many of the phenotypes with significant differences were seed coat pigmentation color and patterning traits. For instance, black and white eye colors and the hilum ring, small eye, Watson, and speckled pigmentation patterns were more common in the WNCA cluster compared to the ESA cluster (see below for a detailed description of these patterns). In contrast, the brown, red, and tan eye colors were more common in the ESA cluster. Additional phenotypes with differential frequencies between the two clusters included several leaflet shapes, with globose and sub-globose leaflet shapes more common in ESA and the sub-hastate shape more common in the WNCA cluster. We also observed that flower pigmentation is more common in the ESA cluster and incidence of target leaf spot was more common in the WNCA cluster.

Taken together, the data suggests that ESA and WNCA cowpea are differentiated across a wide suite of traits spanning plant architecture, pigmentation, and life history.

### Decay of linkage disequilibrium in the cowpea genome

To establish the IITA core collection as a valuable resource for identifying the genetic basis of agronomically relevant traits using GWAS in cowpea, we examined the patterns of genome-wide and chromosome-specific linkage disequilibrium (LD) between SNP markers. Overall, we found that LD decays to R^2^ = 0.20 by 121.5 Kbp on average, with full decay to the genome-wide background by 2.76 Mbp (Figure 4A). We also observed differences in the rate of LD decay between the ESA and WNCA subpopulation clusters, with more rapid decay in the WNCA cluster (R^2^ = 0.20 by 136 Kbp and 89 Kbp, genome-wide background by 3.57 Mbp and 2.97 Mbp, respectively).

The rate of LD decay also varied between chromosomes. LD decay to R^2^ = 0.20 ranged from 68 Kbp on Vu08 to 178 Kbp on Vu01 and decay to chromosome-specific background levels of LD ranged from 721 Kbp on Vu04 to 2.48 Mbp on Vu01 (Figure 4B, Figure S7). As expected given the genome-wide data, the rate of LD decay was more rapid across nearly all chromosomes in the WNCA cluster compared to the ESA cluster.

The resolution of genetic mapping in GWAS is determined by the recombination rate, which determines LD decay, and the density of genetic markers near an associated variant. In regions with dense markers and high recombination (i.e. rapid LD decay), there is a higher likelihood of pinpointing an associated locus within a narrow genomic window. Using our measures of LD decay, we estimate that an associated locus will contain ~25 genes and ~60 SNPs on the array on average. This estimate is based on the average distance of 5,611 bp between adjacent genes, an average gene length of 3,843 bp, and average distance of 3,997 bp between adjacent SNPs as measured in the reference genome (Muñoz-Amatriaín *et al*. 2017; Lonardi *et al*. 2019). However, we acknowledge that these estimates should be considered to be coarse due to the non-uniform distribution of genes and SNPs and variation in the local recombination rate within the genome.

### The genetic architecture of seed pigmentation patterning

To demonstrate the utility of the genotyped IITA core collection to map the genetic basis of agronomically important traits, we focused on studying the genetic basis of seed pigmentation. Variation in the color and pattern of cowpea seed pigmentation is a particularly conspicuous example of the rich phenotypic diversity of the IITA collection. Cowpea seeds range from having no pigmentation (called “no color”) to complete pigmentation throughout the seed coat (so-called “full coat”). Between these extremes, various pigmentation patterns exist, traditionally classified qualitatively. The “small eye” or “Eye 1” pattern displays pigmentation in a narrow margin around the seed hilum with an indistinct margin, while the “large eye” or “Eye 2” pattern displays pigmentation encompassing the hilum with a wide, distinct margin and covering a significant portion of the ventral surface (Herniter *et al*. 2019). The “Watson” pattern, named after a corresponding variety, features pigmentation surrounding the hilum and extending to the micropylar end of the seed, with an unclear boundary (Spillman 1911). Similarly, the “Holstein” pattern, also named after a variety possessing the pattern, exhibits solid pigmentation over most of the seed coat, with patches of color on the remaining portion of the seed. In addition to pigmentation patterning, seed coat color exhibits extraordinary variation across a spectrum of colors including white/cream, tan/buff, brown, red, maroon, and purple/black (Harland 1919, 1920; Spillman and Sando 1930; Saunders 1960; Drabo *et al*. 1988).

Seed coat pigmentation patterning in cowpea has been described as being controlled by a three locus system consisting of the color factor (*C* locus), Watson factor (*W* locus), and Holstein factor (*H* locus) (Spillman and Sando 1930; Saunders 1960; Drabo *et al*. 1988). The *C* locus is epistatic to the other two loci and determines if pigmentation is present throughout the seed coat or just restricted to the eye. The *W* and *H* loci meanwhile act as modifiers and determine whether the pattern of pigmentation in the seed coat develops into the Watson, Holstein, or full coat patterns. Previous work based on QTL mapping in biparental populations has proposed *Vigun07g110700* as a candidate gene for the *C* locus, *Vigun09g13900* for the *W* locus, and *Vigun10g163900* for the *H* locus (Herniter *et al*. 2019).

We first sought to confirm the previously proposed candidate loci for the *W*, *H*, and *C* factors using genome-wide association study in the IITA core collection. Our mapping results for pigmentation patterning yielded three distinct associated loci: one locus on Vu09 and two loci on Vu10 (Figure 5, Table S9). The locus on Vu09 was associated with all five pigmentation patterns, spanned 1.38 Mbp (Vu09:30,035,583-30,035,583), and contained 116 genes (Figure 5F). We identified 2_01960 as the tag SNP for this locus and found that the small eye, large eye, and Holstein pigmentation patterns were associated with the A allele (MAF = 0.146) at this locus while the Watson and full coat patterns were associated with the major frequency G allele. Examination of the genes in this significant interval found that *Vigun09g13990*, a WD-repeat gene proposed as a candidate gene for the *W* locus (Herniter *et al*. 2019), is present in the region between the tag SNP and a linked upstream significant SNP. Furthermore, the candidate gene is surrounded by other significant variants that are not linked with the tag SNP, suggesting the presence of additional associated haplotypes segregating in this region.

The first locus on Vu10 was associated with the small eye, Watson, and full coat pigmentation patterns. The tag SNP for this locus is 2_31919, with the C allele (MAF = 0.221) associated with the small eye and Watson patterns and the major frequency T allele associated only with the full coat pattern. The locus spans 12 KB (Vu10:38,454,820-38,467,161) and overlaps with *Vigun10g165400*, an R2-R3 MYB transcription factor (Figure 5G). R2-R3 MYB genes are reported to have a role in transcriptional regulation of flavonoid biosynthesis as part of MYB-bHLH-WDR complexes (Xu *et al*. 2015).

The genome-wide association study for the Holstein pattern yielded two peaks on Vu10 that are several megabases apart. The first peak has tag SNP 2_15395, which is moderately linked (R^2^ = 0.346) to 2_23640, a SNP that is ~52 KB upstream of *Vigun10g163900*, an E3 ubiquitin ligase previously proposed as a candidate gene for the *H* locus (Herniter *et al*. 2019) (Figure 5H). There are also several SNPs surrounding the gene that did not reach significance per the conservative Bonferroni threshold but are close to the threshold and would have likely been significant with increased power. The second peak spans 189 KB (Vu10:41100865-41290623) and contains 23 genes, but does not yield a clear candidate gene.

Our results confirm that previously reported candidate genes for the *W* locus and *H* locus could be mapped with the Core Collection GWAS panel. Although we were unable to map the *C* locus on Vu07 in the five main seed pigmentation patterns, the locus was mapped in the GWA mapping for hilum ring and eye absent patterns (Figure S8).

### The genetic architecture of seed color

In cowpea, seed coat color varies on a continuous spectrum with several colors typically described qualitatively such as white/cream, tan/buff, brown, red, blue, purple, and black. Previous genetic studies of the segregation of seed coat color phenotypes revealed that coat color is inherited independently of pigmentation pattern and is likely controlled by at least five unlinked major genes (Saunders 1960; Drabo *et al*. 1988). To date, only the locus controlling black seed color, the *Bl* locus, has been identified (Herniter *et al*. 2018). To attempt to identify the loci associated with additional seed coat colors, we mapped the genetic basis of the red, tan, black, blue, and white seed coat colors using GWAS in the Core Collection (Figure 6). In total, we found 4 major peaks associated with seed coat color: one each on Vu03 (red), Vu05 (black), Vu06 (blue), and Vu10 (No color/white).

We first sought to confirm if the Vu05 locus previously proposed to be associated with black seed coat color (Herniter *et al*. 2018) could be mapped via GWAS in the core collection. The GWAS for black seed coat yielded a major 121 KB peak on Vu05 (Vu05:3,104,538-3,225,877), spanning 13 genes (Figure 6A, F). The locus was also associated with the tan seed coat color (Figure 6C). The minor C allele at tag SNP 2_19309 (MAF = 0.463) was associated with the black seed coat phenotype while the major T allele was associated with the tan seed coat phenotype. Among the genes in the region, we found *Vigun05g039500*, a R2-R3 MYB transcription factor previously identified as a candidate gene for the *Bl* locus, which is associated with black seed coat and purple pod tip (Herniter *et al*. 2018).

The peak on Vu03 is associated with both the red and tan seed colors (Figure 6B-C, G), spans 274 KB (Vu03:10933603-11208404) and contains 25 genes. The T allele (MAF = 0.102) at the tag SNP 2_20787 is associated with red seed color, while the major frequency G allele is associated with the tan color. The locus maps to a pair of anthocyanin reductase (ANR) genes: *Vigun03g118700/Vigun03g118800*, with the tag SNP a predicted missense mutation in the former gene. ANR acts in the biosynthesis of flavan-3-ols and proanthocyanidins (condensed tannins) in plants (Xie *et al*. 2003). Mutations in ANR in *Arabidopsis thaliana* have been shown to result in the accumulation of anthocyanins in the seed coat (Devic *et al*. 1999), making this locus a compelling candidate for the red seed color phenotype.

The GWAS for blue seed color resulted in a ~155 KB peak on Vu06 (Vu06:13491073-13645944) that spans 5 genes (Figure 6D). SNP 2_22603, which is strongly linked to the tag SNP and lies directly upstream, is a predicted synonymous or intron mutation within the gene *Vigun06g030900* (Figure 6H). This gene is a homolog of *AT3G59030* (*TRANSPARENT TESTA12*) which has been shown to be required for sequestration of flavonoids in the seed coat epithelium tissue of *A. thaliana (Debeaujon et al. 2001)*.

Our attempt to map the genetic basis of white/cream seed color yielded one of the peaks on Vu10 (2_31919) observed in seed coat patterning GWAS (Figure 5). Additionally, we did not observe any significant SNPs in the GWA mapping of brown seed coat color, likely due the fact that this phenotype can be heat-induced (Pottorff *et al*. 2014). In total, our GWAS was able to map a previously identified locus for the black seed coat color and propose novel loci associated with the red, tan, and blue seed coat colors.

### Mapping of additional phenotypes

Including the seed coat pigmentation pattern and color phenotypes discussed previously, we attempted to map the genetic basis of 71 distinct phenotypes in total. Altogether, 42 of the genome-wide association studies had one or more significant SNPs (range 1 to 106 SNPs, median 8). Of the 748 SNPs that were associated with variation in a phenotype, 698 were associated with variation in only one phenotype and 50 SNPs were associated with more than one phenotype (Table S9).

We found numerous interesting association peaks with clear underlying candidate genes. For instance, GWAS for presence of flower pigmentation had a significant SNP adjacent to *Vigun07g148800*, an E3 ubiquitin ligase. Although E3 ubiquitin-ligases are implicated in numerous biological processes, they have been reported to be negative regulators of the flavonoid biosynthesis pathway transcriptional activation complex (Shin *et al*. 2015; Herniter *et al*. 2019). We also found candidate genes associated with disease resistance. GWAS for incidence of rust yielded two peaks with clear candidates: a significant SNP on Vu02 in a 12-oxophytodienoate reductase (*Vigun02g192500*) involved in the Jasmonic acid biosynthesis pathway and a very strong peak on Vu09 spanning 90 KB (Vu09:2156441-2246970) containing three TIR-NBS-LRR family disease resistance genes with substantial sequence similarity: *Vigun09g027450*, *Vigun09g027600* (DSC1), and *Vigun09g027700* (suppressor of npr1-1). In sum, our analyses demonstrate the utility of the genotyped IITA Cowpea Core Collection for discovering variants associated with phenotypic variation in cowpea.

## Discussion

The purpose of this study was to utilize the recently reported SNP genotyping data to examine population structure and the potential for genome-wide association mapping in the IITA Cowpea Core Collection. The accessions in this collection are representative of the genotypic and phenotypic diversity of extant cultivated cowpea germplasm. To our knowledge, the dataset constitutes the largest cohort of cowpea that have been genotyped genome-wide with a single technology and we anticipate that the resources linked to this work will be of considerable utility to the cowpea research and breeding community.

### Genetic relationships in global cowpea

Our analysis of population structure revealed that global cowpea is broadly genetically differentiated into two major subpopulations defined by geography, with separation between germplasm collected in Eastern and Southern Africa (ESA cluster) from germplasm collected in West, Central, and North Africa (WNCA cluster). The genetic differentiation between germplasm from West Africa and East Africa has been observed previously in smaller subsets of globally collected cowpea germplasm (Huynh *et al*. 2013; Xiong *et al*. 2016; Muñoz-Amatriaín *et al*. 2017, 2021; Fatokun *et al*. 2018; Herniter *et al*. 2020), and our observations confirm that this is the major organizing principle of cowpea diversity. However, we did observe that between 9-12% (depending on the number of subpopulations considered) of accessions collected in West Africa were assigned to the ESA cluster (cluster 1). While this pattern is consistent with the proposed domestication of cowpea in West Africa, our data cannot rule out the possibility of a separate domestication center in East Africa. Additional study of wild cowpea in both West Africa and East Africa could be useful for reconstructing the mode and tempo of domestication of this crop.

Although the IITA Cowpea Core Collection primarily consists of traditional cultivars from West Africa, the population structure analyses did provide some insight into the dispersal of cowpea germplasm around the globe. The majority of germplasm from Latin America and the Caribbean, North America, and Oceania clustered within the ESA cluster, suggesting dispersal to these locales from Eastern and Southern Africa. In contrast, germplasm collected from East Asia, Melanesia, and Southeast Asia was mostly assigned to cluster 3 and thus likely represents migrants dispersed out of North Africa, perhaps through the Sabaean Lane, an ancient trade route connecting East Africa with modern-day India (Murdock 1959; Herniter *et al*. 2020). Further, our analyses indicated a limited role of cluster 2 (West Africa specific) ancestry in the dispersal of cowpea out of Africa, with only a few accessions, all present in Latin America and the Caribbean, assigned to this cluster when considering more than 2 subpopulations. Our analysis of phenotypic differences between the ESA and WNCA clusters suggested that ESA germplasm is associated with earlier flowering, suggesting that this phenotype was important for the worldwide dispersal of cowpea.

Compared to other surveys of global population structure in cowpea, our analysis yielded fewer “optimal” clusters. We attribute this result to our use of the Evanno method (Evanno *et al*. 2005) to decide the number of subpopulations, which has been shown previously to be biased to select K = 2 (Janes *et al*. 2017). Therefore, we considered up to two additional subpopulations, identifying a cluster of accessions collected from the Nile Delta in Northern Egypt in close proximity to the Mediterranean Sea and a cluster of early flowering accessions in West Africa, both of which were also discovered in the UCR Minicore, a mini-core subset of a cowpea diversity collection held at the University of California, Riverside (Muñoz-Amatriaín *et al*. 2021). In contrast to previous studies, we did not observe subpopulation clusters consisting of only breeding lines or clusters consisting solely of germplasm collected outside of Africa. It is possible that these signals are also present in the data and would have been resolved if we had improved sampling from continents beside Africa, if we had considered additional subpopulations, or if we had conducted the analysis omitting the traditional cultivars from West Africa that constitute the bulk of the collection. Our analysis also did not identify the cluster of yardlong bean (subspecies sesquipedalis) previously identified in other collections of global germplasm (Muñoz-Amatriaín *et al*. 2021) as the IITA Cowpea Core Collection only has three accessions known to be from this group.

In addition to the patterns of population structure, we found that a substantial number of accessions are nearly identical to one or more other accessions in the Core Collection. Our analysis indicated that the majority of clusters of identical accessions consisted of lines collected across geopolitical boundaries, with many clusters containing collections transcending continents. A primary objective when selecting accessions for the core collection was to ensure that the chosen lines were representative of the geographic and phenotypic variation present in the full IITA cowpea collection (Mahalakshmi *et al*. 2007). Consequently, the occurrence of dispersed identical lines within the core collection may be a result of sampling across geographic gradients (i.e. sampling an accession that had been dispersed across a geopolitical border in both the original and dispersed location in an effort to have representatives from all regions where cowpea is found) or due to sampling of lines subject to phenotypic plasticity upon selection of the core collection subset.

### A genome-wide association mapping panel for cowpea

A primary objective of this study was to determine whether or not the IITA Cowpea Core Collection constitutes a suitable panel for genome-wide association mapping in domesticated cowpea. The rate of decay of linkage disequilibrium dictates the precision with which significant associations can be linked to specific candidate genes. The decay of linkage disequilibrium (LD) in the cowpea core collection is slow when compared to other plant systems (1.5 KB in maize (Remington *et al*. 2001) and 10 KB in *A. thaliana (Kim et al. 2007)*), which is consistent with observations in smaller previously genotyped collections (Muñoz-Amatriaín *et al*. 2021; Sodedji *et al*. 2021). The slow decay of LD may be a feature common to legumes as similar rates have been reported in cultivated soybean (Zhou *et al*. 2015). However, the slow decay of LD in cowpea is partly due to strong population structure, as LD decays somewhat more quickly within the WNCA population cluster.

Despite the slow decay of LD in cowpea, we were able to use genome-wide association to identify hundreds of loci associated with variation in dozens of traits. We note that the phenotype values used for our analyses were collected across many field studies spanning years and locations, leaving us unable to control for the contribution of environmental variance to phenotypic variance. We anticipate that GWAS of phenotypes collected in a single experiment will have even greater success in identifying the genetic basis of trait variation using the IITA Cowpea Core Collection.

### The genetic basis of seed coat pigmentation in cowpea

Our efforts to map the genetic basis of seed pigmentation patterning using GWAS yielded several candidate loci that were previously discovered using QTL mapping in F2 and advanced RIL populations (Herniter *et al*. 2019). The consistency between the loci mapped as part of our work and previous reports suggests that the loci proposed previously likely regulate seed pigmentation pattern more generally in global cowpea and support the three locus model for the genetic control of seed pigmentation patterning (Spillman and Sando 1930; Saunders 1960; Drabo *et al*. 1988).

Through GWAS, we were able to map a second locus on Vu10 that likely contributes to seed pigmentation patterning (Figure 5G). For this locus, we propose *Vigun10g165400*, a R2-R3 MYB transcription factor, as a candidate gene. Analysis of patterns of linkage disequilibrium at this locus suggests that this locus is distinct from the candidate gene at the *H* locus despite being in close proximity (~110 KB downstream). This locus has been observed in a previous attempt to map the genetic basis of seed pigmentation patterning (Herniter *et al*. 2019), but was disqualified due to not being observed in the associated haplotype blocks of all the tested F2 populations. Our results support the hypothesis that multiple genes in close proximity of Vu10 likely regulate seed pigmentation patterning. Further work is needed to determine the molecular basis of this trait and to demonstrate the efficacy of the proposed candidates.

Seed coat color has been studied for over 100 years in cowpea (Harland 1919, 1920) and the existence of at least five genes affecting this trait have been proposed (Saunders 1960; Drabo *et al*. 1988). However, the full genetic control of seed coat color has yet to be elucidated. Here, we propose that the *R* locus controlling red seed color is an anthocyanin reductase (ANR), either *Vigun03g118700* or *Vigun03g118800*. Although both genes lie under the associated peak, the tag SNP is a predicted missense variant in *Vigun03g118700*, which is also expressed at a high level during early seed development in the reference cultivar IT97K-499-35, which has a black-eyed phenotype. In contrast, *Vigun03g118800* is expressed at a low level across many tissues and developmental timepoints (Yao *et al*. 2016). Thus, we consider *Vigun03g118700* to be the likely causal locus for the red seed coat color phenotype.

ANR is active in the flavonoid biosynthesis pathway, where it competes with 3-O-glucosyltransferase to convert cyanidin into condensed tannins or anthocyanins, respectively (Kovinich *et al*. 2012). Mutations in ANR have been shown to result in the rapid accumulation of anthocyanins during early seed development in *Arabidopsis thaliana* (Devic *et al*. 1999) and knockdown of ANR in soybean yields a red seed coat phenotype (Kovinich *et al*. 2012). Our results suggest that a non-functional ANR gene is responsible for the red seed coat phenotype in cowpea.

We also propose *Vigun06g030900*, a homolog of *TRANSPARENT TESTA12 (TT12)*, as a candidate gene for the blue seed color phenotype. *TT12* is a MATE transporter which mediates anthocyanin deposition in *A. thaliana* seeds (Debeaujon *et al*. 2001; Marinova *et al*. 2007). It remains to be seen whether blue seed color might be driven by changes in activity or regulation of *Vigun06g030900*.

A full genetic model of seed pigmentation patterning and color in cowpea based on our GWAS is presented in Table 1.

**Table 1.**
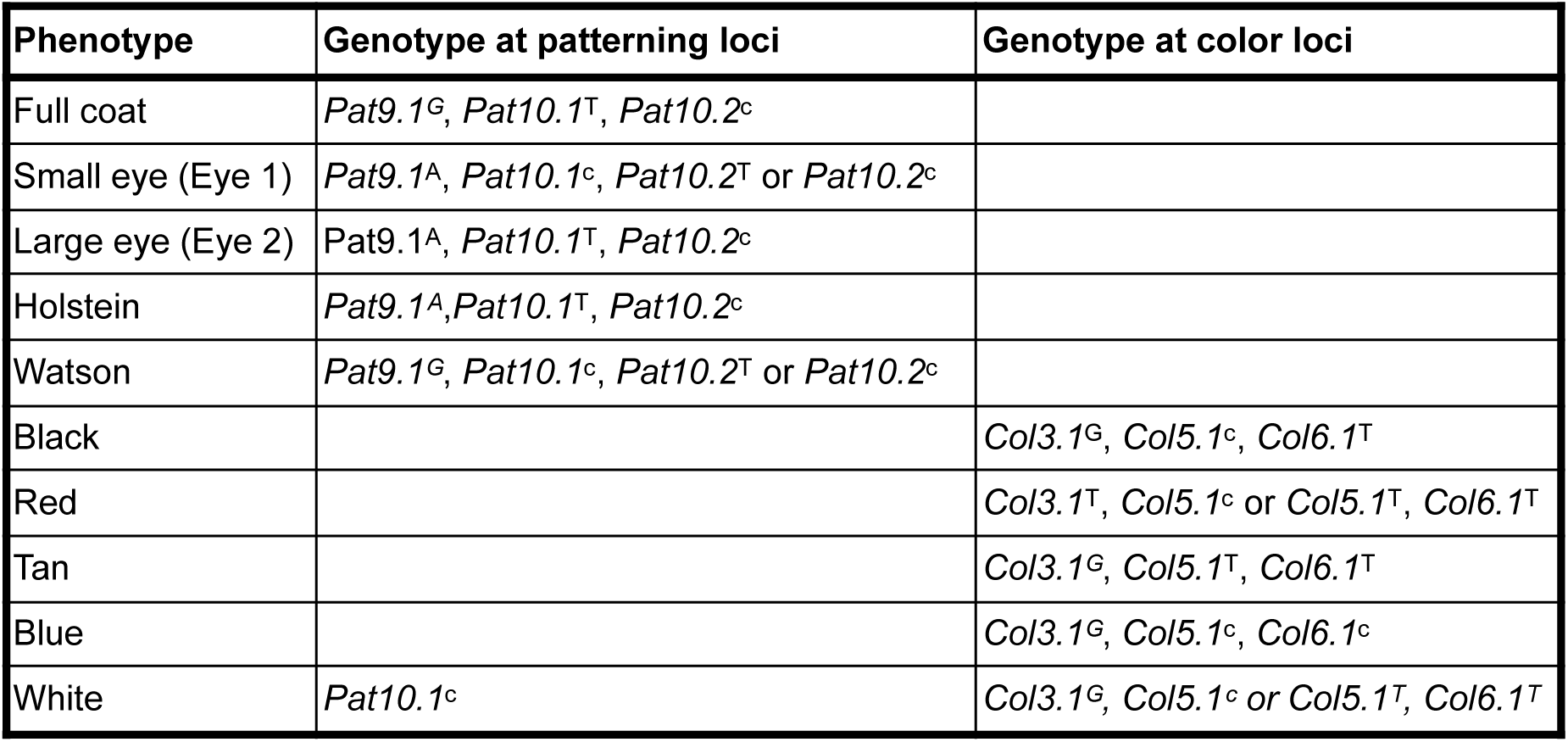
Genetic model of seed pigmentation pattern and color. Proposed genotypes at pigmentation patterning and color quantitative trait loci. QTL correspond to SNPs as follows: *Pat9.1* = 2_01960, *Pat10.1* = 2_31919, *Pat10.2* = 2_15395, *Col3.1* = 2_20787, *Col5.1* = 2_19309, Col6.1 = 2_22603.

### Future directions

The genotyped IITA Cowpea Core Collection represents a valuable tool for dissecting the genetic basis of agronomic traits in cowpea. We anticipate that the resources produced by this work will be useful for marker-assisted breeding and will facilitate the production of new cultivars in response to consumer preference and changing environmental conditions. To facilitate the latter aim, this collection could be used to identify climate-associated alleles, as many of the traditional cultivars have known collection locations. This knowledge could be especially useful for ensuring the resilience of cowpea into the future.

## Materials and Methods

### Accession collection sites

Collection sites for each accession were provided by the IITA (IITA 2023). World regions are described according to the United Nations M49 standard (United Nations Statistics Division).

### Genotyping and variant filtering

Accessions were genotyped using the Illumina Cowpea iSelect Consortium Array which assays nearly 50,000 SNPs (Muñoz-Amatriaín *et al*. 2017) as described in (Close *et al*. 2023). All alleles reported in this work belong to the Watson strand.

SNPs with greater than 95% of missing calls across samples were filtered first, followed by filtering of samples with greater than 50% missing calls or more than 20% heterozygous calls across sites. Near identical lines were identified by calculating pairwise Hamming distance between samples with plink 1.9 (Chang *et al*. 2015), applying hierarchical clustering to the distance matrix, and defining clusters of accessions using a static tree cut at height of 500. We randomly selected one sample from each cluster to retain for subsequent analysis.

From the filtered SNP set, we also produced a set of SNPs in approximate linkage equilibrium by first subsetting to SNPs with minor allele frequency greater than or equal to 0.01 with no more than 5% missing calls. The filtered SNPs were then pruned pairwise in 50 SNP windows with a step size of 10 and an R^2^ threshold of 0.20. All SNP filtering and pruning was done using plink 1.9 (Chang *et al*. 2015). The pruned set, composed of 4,245 SNPs, was used for all population structure analyses.

### Variant effect prediction

The predicted effect of variants were determined using Ensembl Variant Effect Predictor v. 95.0 (McLaren *et al*. 2016) using the *V. unguiculata* v. 1.0 reference genome assembly and v. 1.2 of the genome annotation (Lonardi *et al*. 2019).

### Genetic diversity statistics

Allele frequencies and pairwise genetic distances were calculated using plink v.1.9 (Chang *et al*. 2015). Expected heterozygosity was calculated according to the following formula:

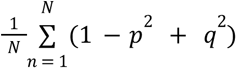, where N is the total number of sites and p and q are the frequencies of the major and minor frequency alleles at each site, respectively.

### Population structure

Population structure was analyzed using ADMIXTURE 1.3.0 (Alexander *et al*. 2009; Alexander and Lange 2011) for K = 1 to 10 with 10 replicates per K. The most likely K value was determined using the Evanno method (Evanno *et al*. 2005). ADMIXTURE runs were post-processed and visualized using the pophelper 2.3.1 R package (Francis 2017). Samples were assigned to the group contributing 70% or more of the sample’s estimated ancestry fraction. All other samples were considered to be unassigned.

Principal component analysis on allele frequencies was done using the adegenet v. 2.1.5 (Jombart 2008) R package. The mean allele frequency at a locus was substituted for missing genotype calls.

### Phenotypic analyses

The IITA Cowpea Characterization and Cowpea Evaluation phenotypic datasets were downloaded from the International Institute of Tropical Agriculture (IITA) website (IITA 2023) and consisted of 15 quantitative and 23 qualitative traits. The data were collected as part of the IITA cowpea breeding program across multiple field trials spanning many years and thus represent reference values used for breeding. For the analysis, seed color pigmentation traits were defined according to the majority color of the seed and similar traits were combined as described in Table S6. “Small eye” is synonymous with the “Eye 1” pattern and “Large eye” is synonymous with the Eye 2 pattern described by (Herniter *et al*. 2019). “Self-colored” is described herein as the synonymous “full coat” pattern (Singh and International Institute of Tropical Agriculture 2014). Qualitative traits were coded into binary “case/control” variables using the fastDummies 1.6.3 R package (Kaplan 2020) or considered to be discrete numerical variables (Table S4). Differentiation in phenotype distribution between population structure clusters was tested using Mann Whitney Wilcoxon tests for quantitative phenotypes and Fisher’s Exact test for qualitative phenotypes in R (R Core Team 2013). Two-sided tests were used for all analyses and a Bonferroni adjusted threshold was used to determine significance at α = 0.05.

### Linkage disequilibrium

Pairwise linkage disequilibrium between markers was calculated with plink 1.9 (Chang *et al*. 2015). Since calculation of pairwise LD for all markers in a dataset can be computationally prohibitive, we only calculated LD between markers that were within 10 Mbp of each other. Genome-wide and chromosome-specific background levels of LD were determined by first calculating the mean R^2^ value in 500 bp distance bins and then taking the median value across all bins.

### Genome-wide association mapping

Univariate mixed linear models were fitted with GEMMA 0.98.4 (Zhou and Stephens 2012). The models included a centered relatedness matrix to correct for spurious associations due to population structure. Only sites with minor allele frequency greater than 0.01 and missing in fewer than 5% of samples were tested in this analysis. Continuous quantitative phenotypes were log-transformed prior to analysis. Qualitative phenotypes were encoded in binary as 1 or 0 corresponding to presence or absence of the trait. Bonferroni correction was used to correct p-values for multiple testing and significance was assessed at α = 0.05. Linked significant SNPs (R^2^ > 0.20) within 1 Mbp were considered to define a single peak and the SNP with the lowest p-value per peak was assigned to be the “tag SNP.” Candidate genes were identified by examining the overlaps between significant peaks and loci described in the *Vigna unguiculata* v.1.2 genome annotation (Lonardi *et al*. 2019).

### Gene expression data

The *Vigna unguiculata* Gene Expression Atlas (VuGEA) data (Yao *et al*. 2016) were retrieved from the Legume Information System (Berendzen *et al*. 2021) at https://data.legumeinfo.org.

## Data Availability

The genotyping data has been published previously on Dryad (https://doi.org/10.6086/D19Q37). Information about the IITA Cowpea Collection, including phenotyping data, is available at https://my.iita.org/accession2/collection.jspx?id=1. All other data necessary for reproduction of this work is present within the manuscript, supplemental figures, and supplemental tables.

## Acknowledgements

We thank the staff of the International Institute of Tropical Agriculture (IITA), especially Ousmane Boukar, for support of this project and Jeffrey Ehlers of the Bill and Melinda Gates Foundation for insights regarding the characteristics particular to West African subpopulations. We also thank the members of the Koenig and Seymour labs at University of California, Riverside (UCR) for their feedback on the analyses and Eric Castillo and Sabrina Phengsy for assistance with the seed photography. All computation was performed using the resources and support provided by the High-Performance Computing Center at UCR. This work was supported by a grant to DK from the The International Maize and Wheat Improvement Center (ALLM-007) and by funding provided by the Crop Trust to IITA.

## Supplemental Figures

**Figure S1.**
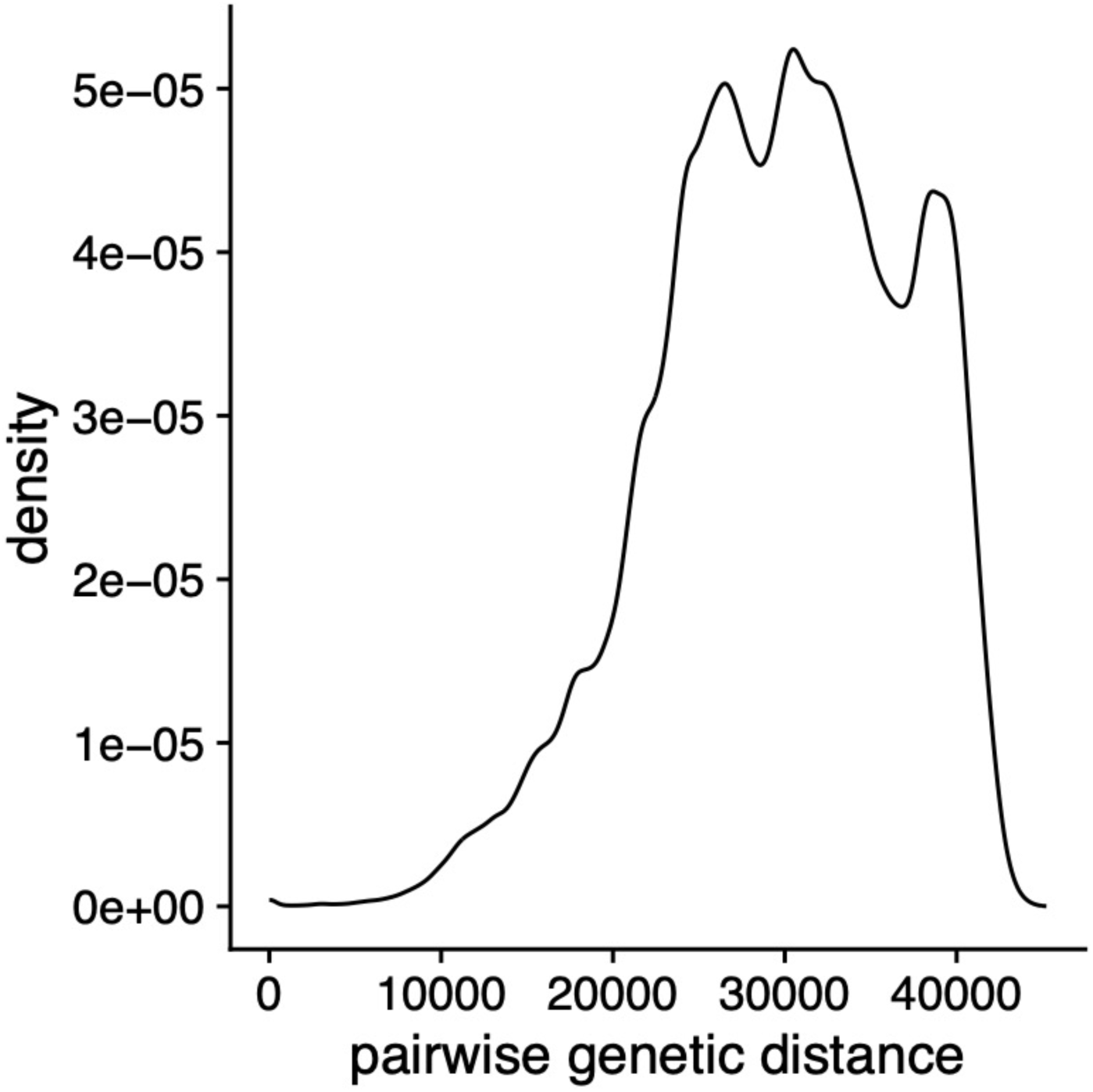
Distribution of pairwise genetic distances between accessions.

**Figure S2.**
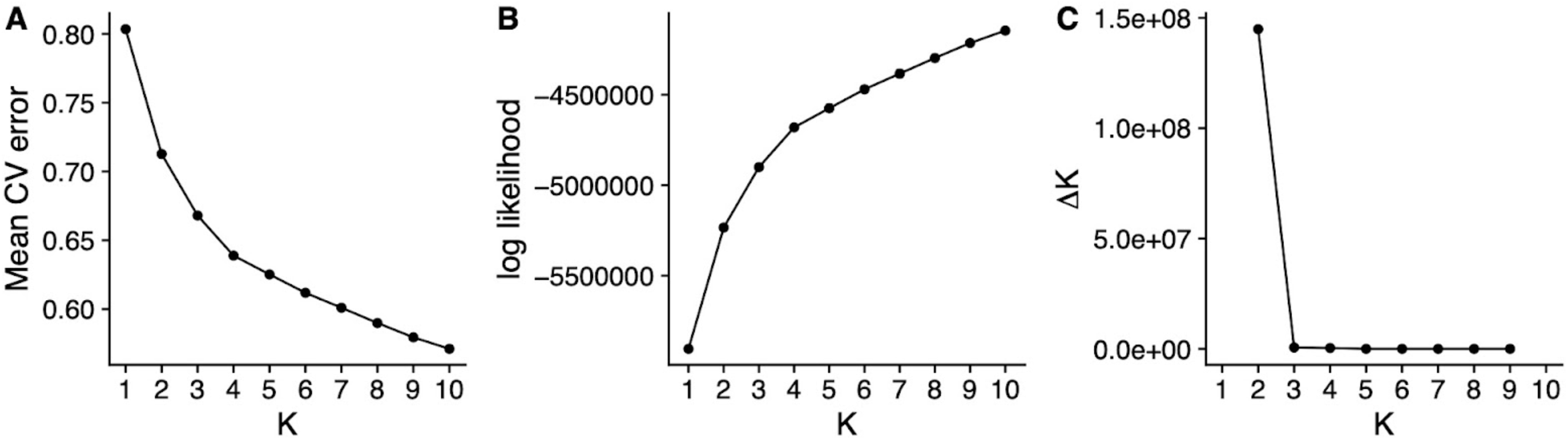
Admixture statistics per K. A) Mean cross-validation error. B) Mean log-likelihood. C) Delta K.

**Figure S3.**
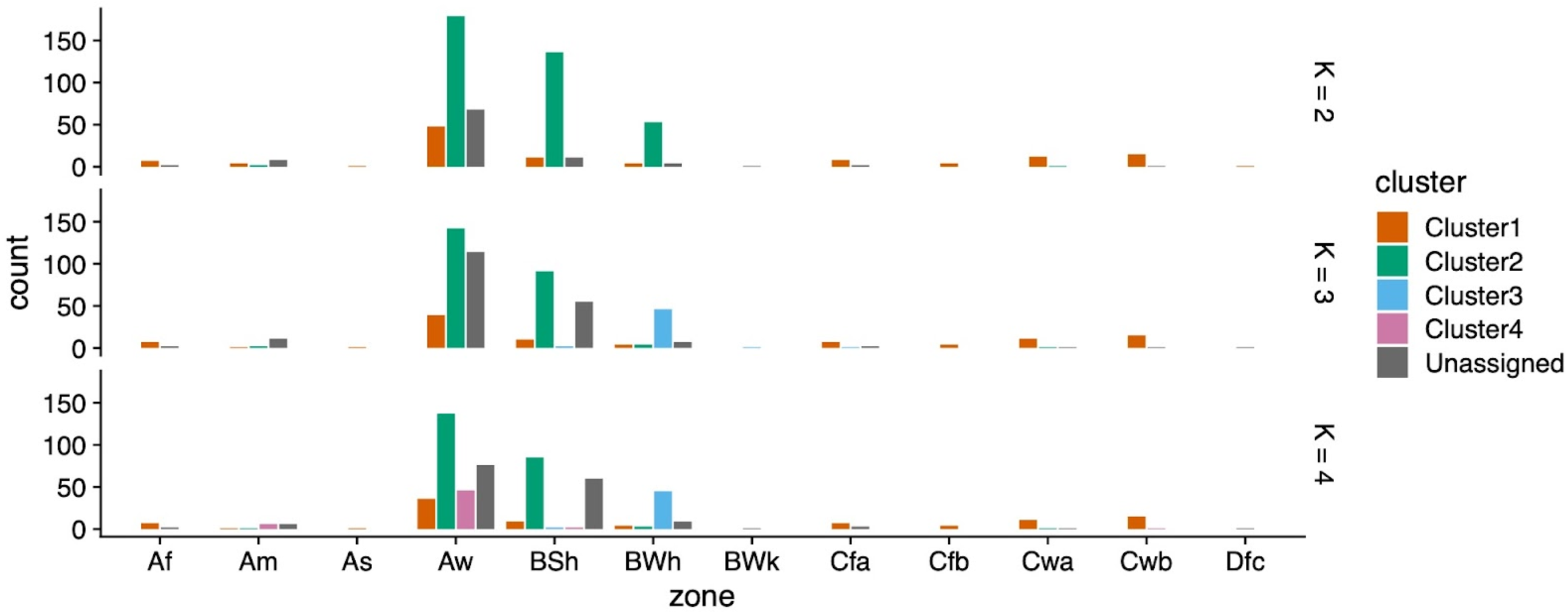
Climate zone distribution by subpopulation cluster for accessions with collection coordinates. Climate designations correspond to the Köppen climate classification.

**Figure S4.**
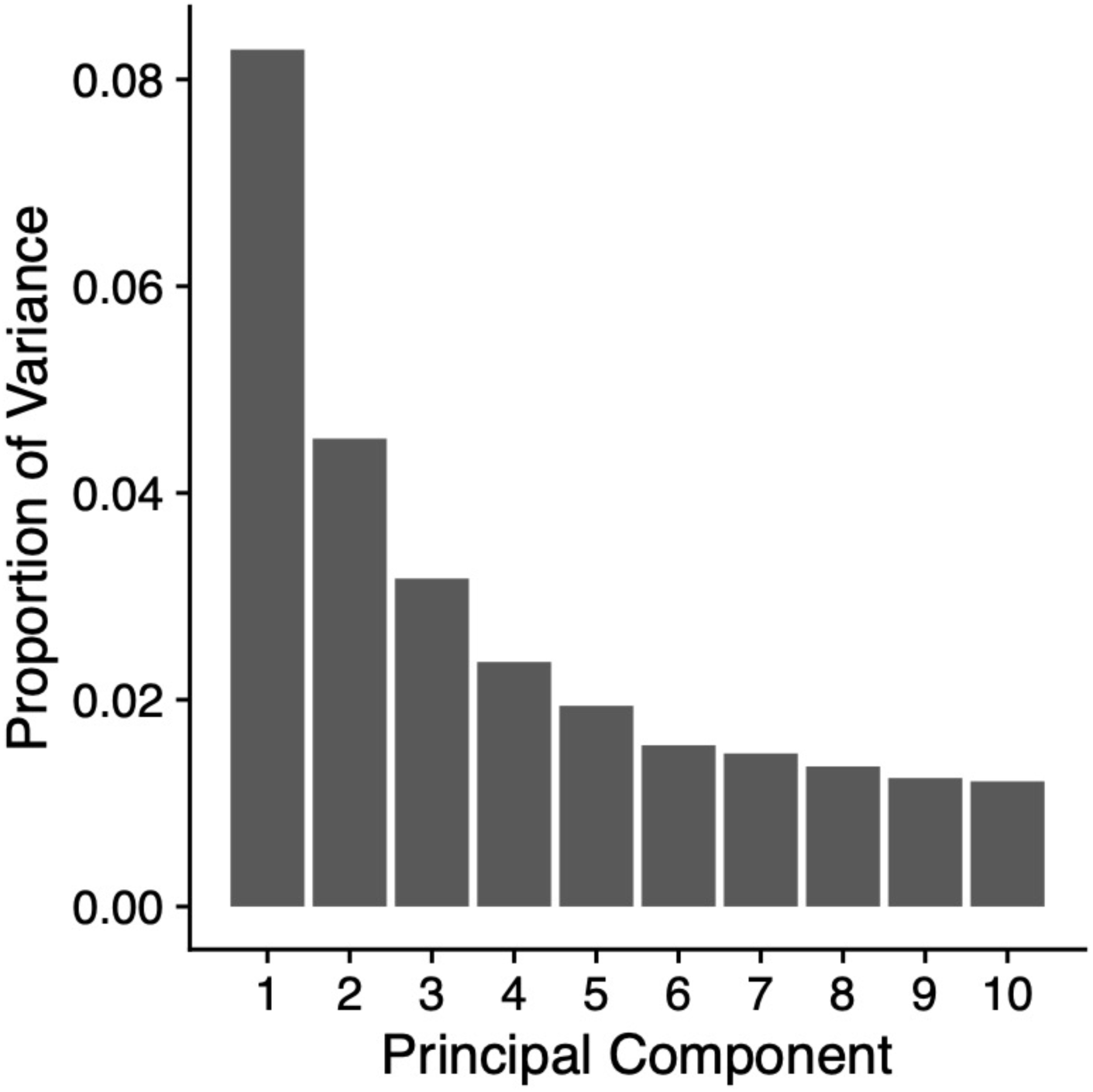
Proportion of variance explained by the first ten principal components.

**Figure S5.**
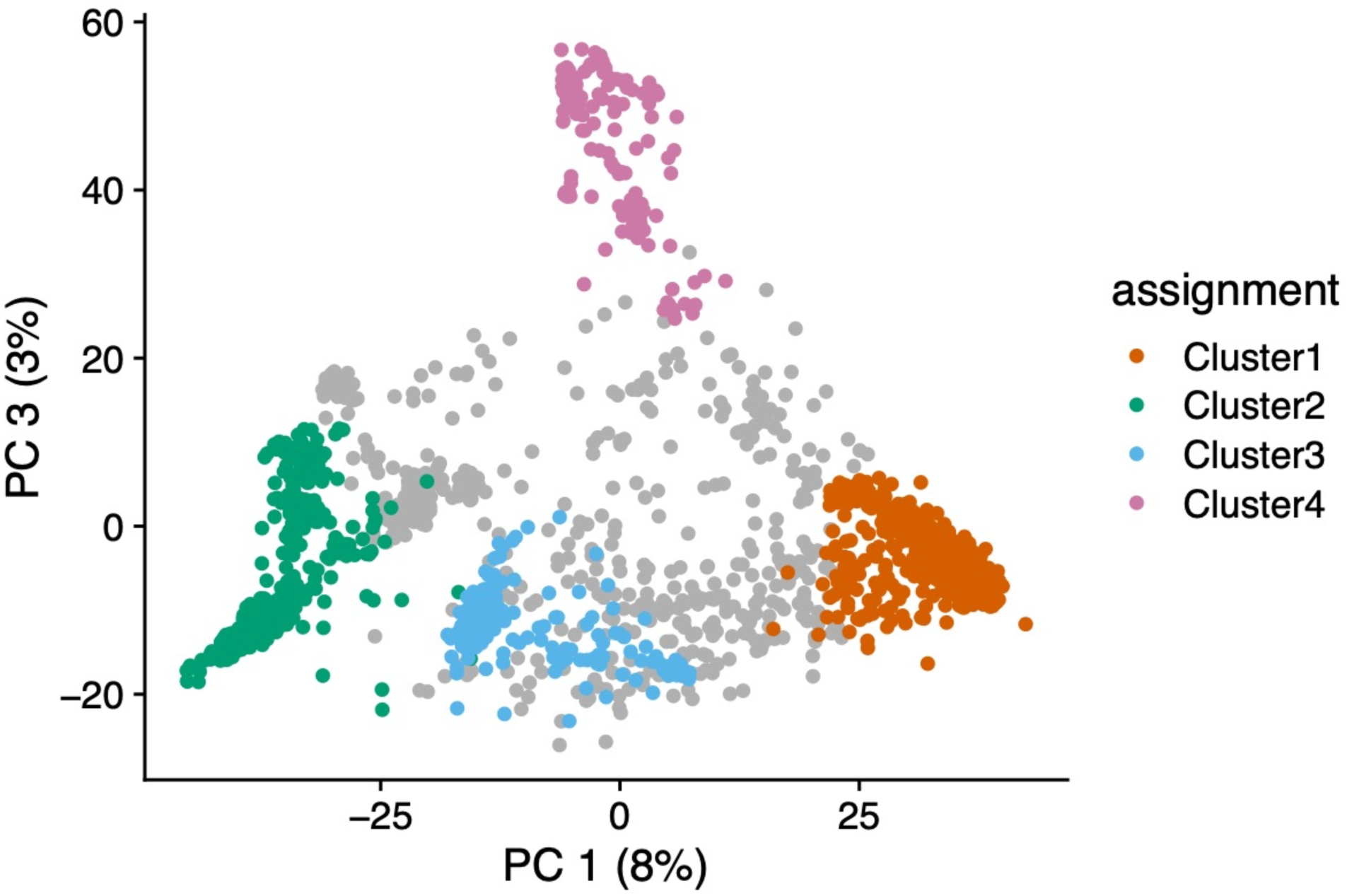
PC 1 vs. PC 3 (K = 4).

**Figure S6.**
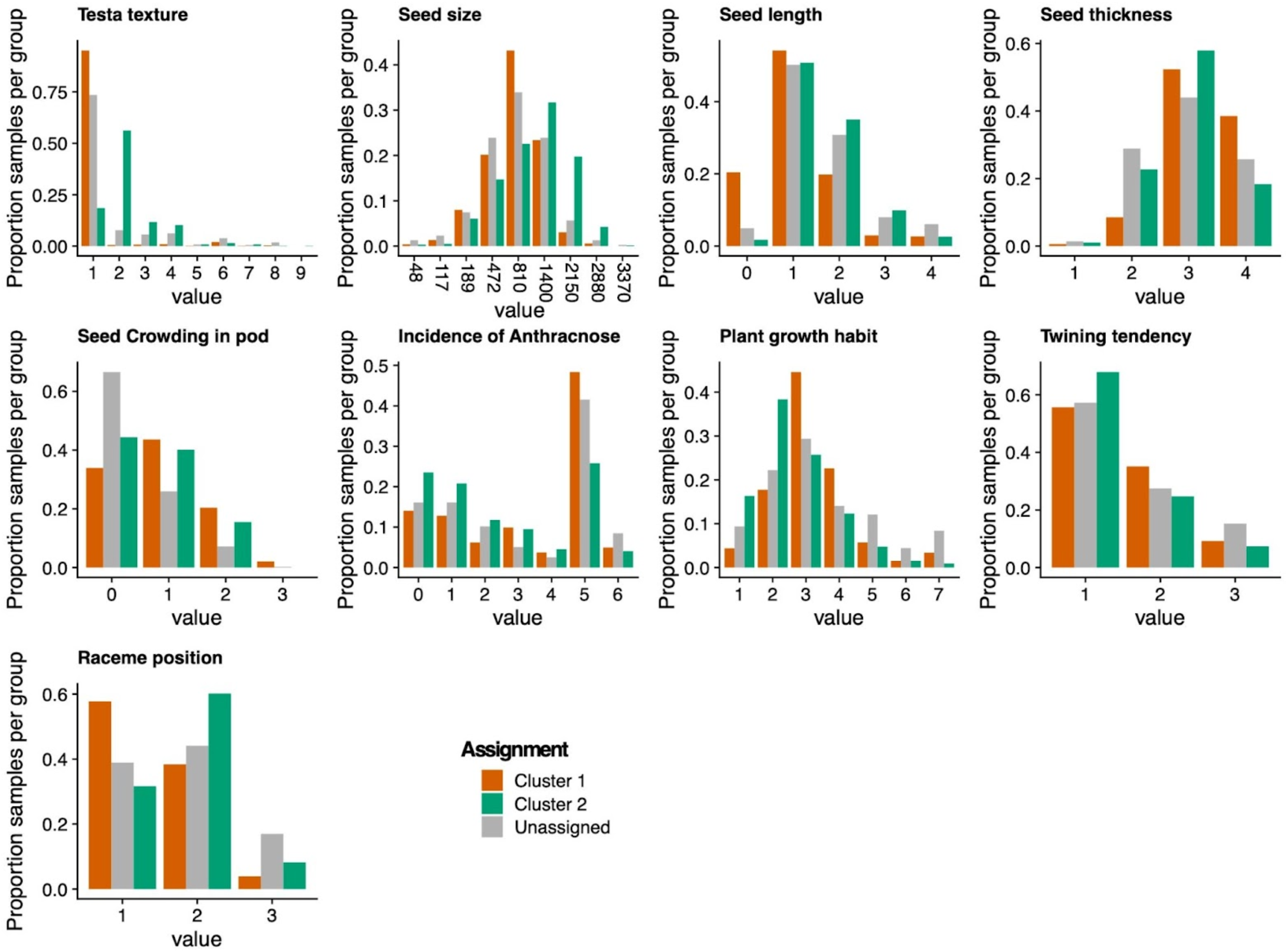
Distributions of discrete phenotypes that vary between Cluster 1 and Cluster 2 (K = 2). All phenotypes shown had a statistically significant difference between the two subpopulation clusters (Mann Whitney Wilcoxon adjusted p-value < 0.05).

**Figure S7.**
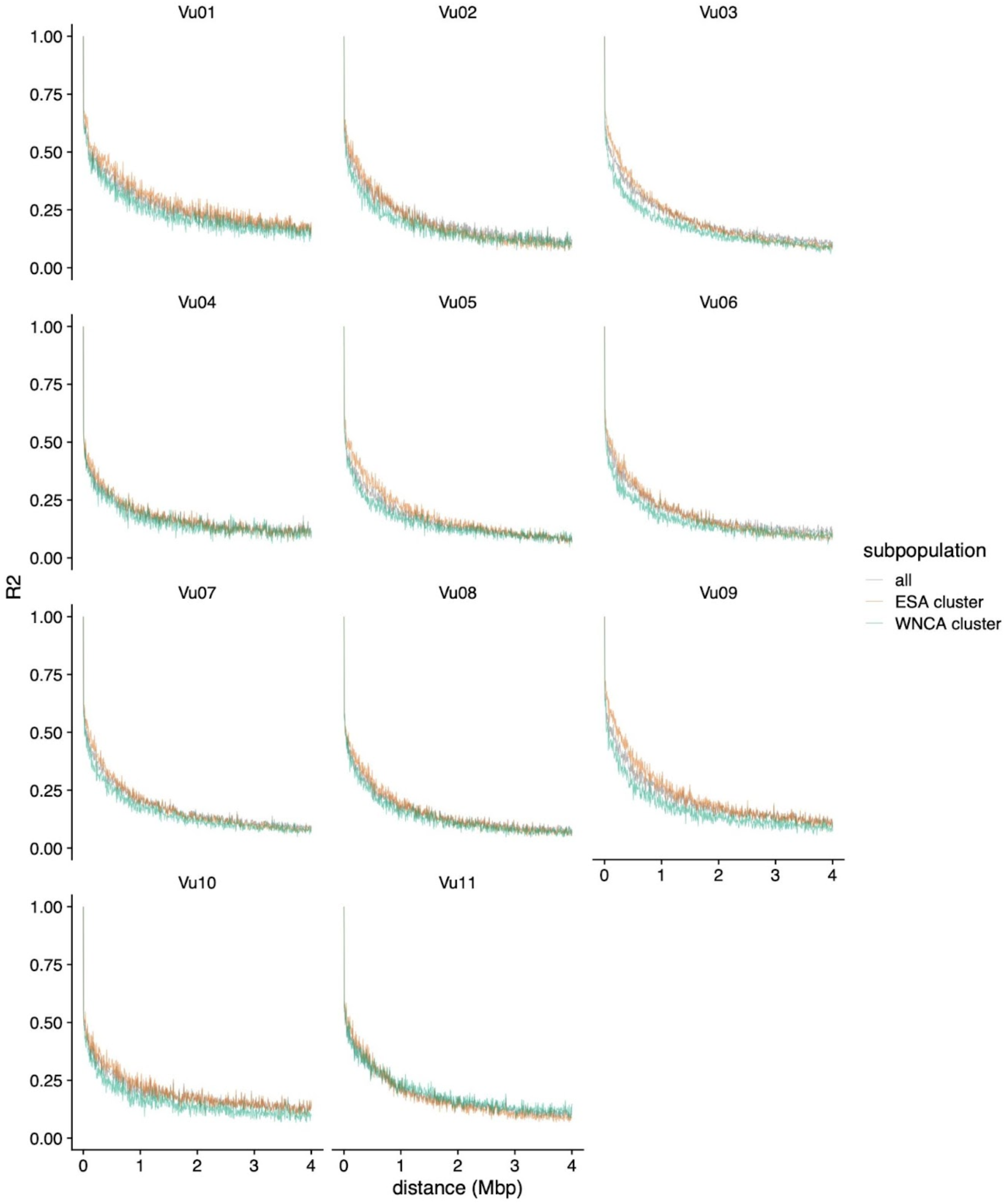
Linkage disequilibrium decay by chromosome.

**Figure S8.**
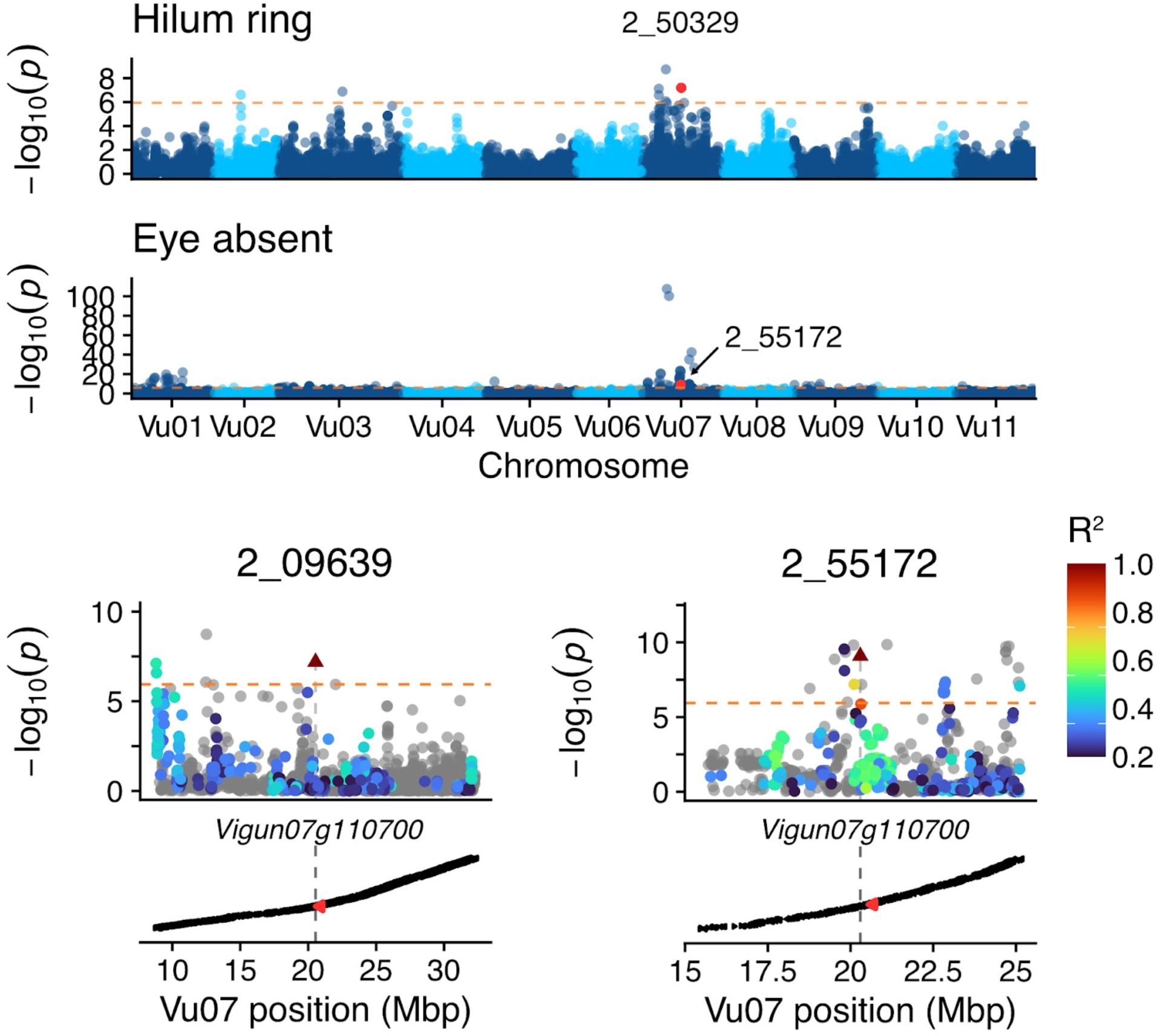
Genome-wide association mapping manhattan plots and focal loci for additional eye pattern phenotypes.

**Figure S9.**
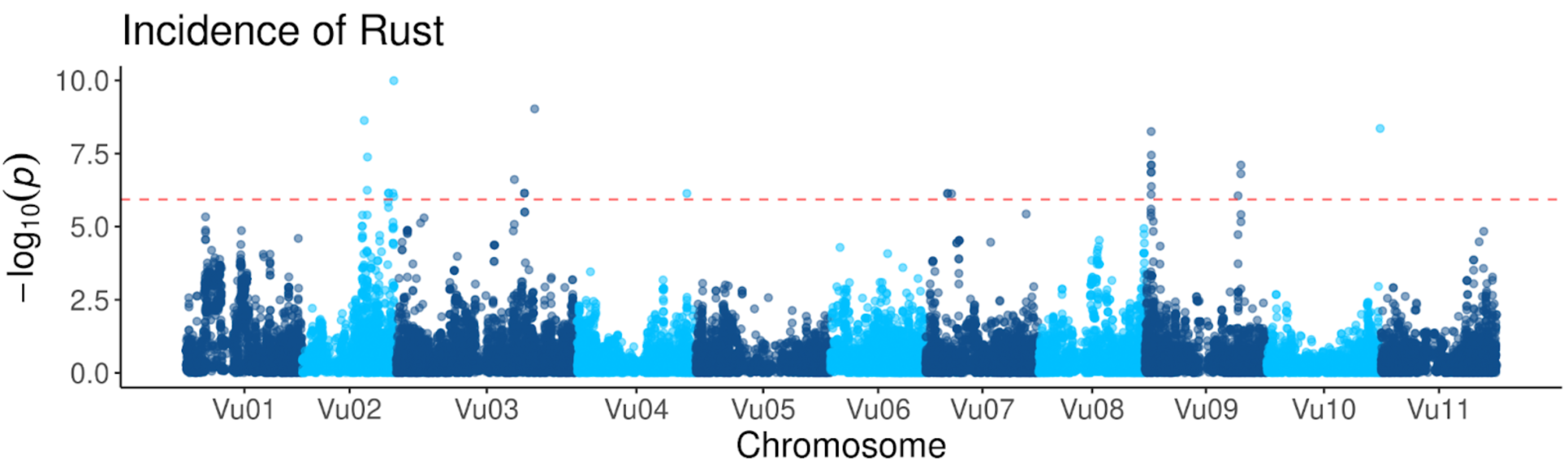
GWAS for rust resistance.

**Figure S10.**
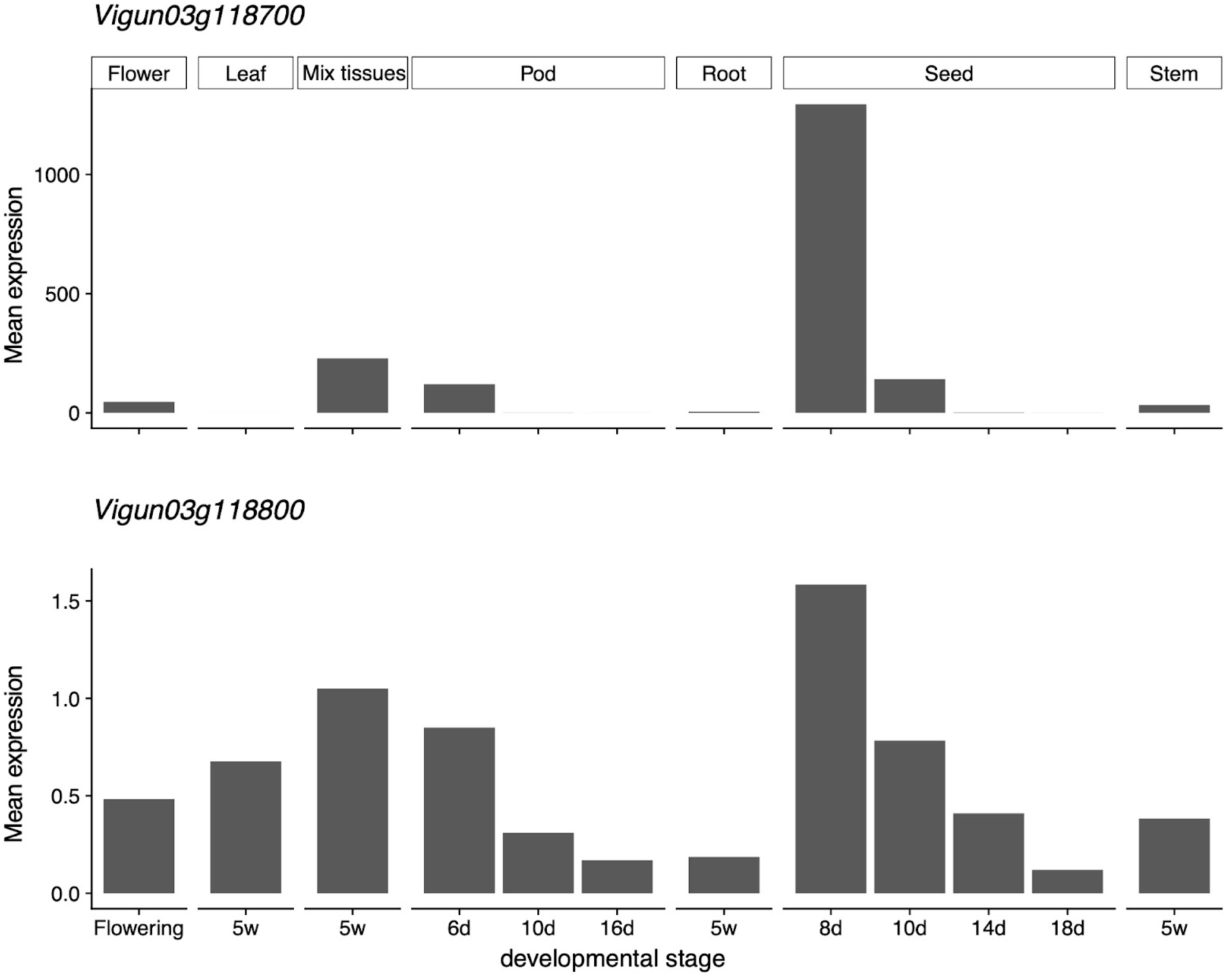
Expression of 2 candidate genes for Tan and Red seed coat color across developmental stages and tissues.

